# The TCR/peptide/MHC complex with a superagonist peptide shows similar interface and reduced flexibility compared to the complex with a self-peptide

**DOI:** 10.1101/588483

**Authors:** Ilaria Salutari, Roland Martin, Amedeo Caflisch

## Abstract

T-cell receptor (TCR) recognition of myelin basic protein (MBP) peptide presented by major histocompatibility complex (MHC) protein HLA-DR2a, one of the MHC class II alleles associated with multiple sclerosis, is highly variable. Interactions in the trimolecular complex between the TCR of MBP83-99-specific T cell clone 3A6 with the MBP-peptide/HLA-DR2a (abbreviated TCR/pMHC) lead to substantially different proliferative responses when comparing the wild-type decapeptide MBP90-99 and a superagonist peptide which differs mainly in the residues that point towards the TCR. Here we investigate the influence of the peptide sequence on the interface and intrinsic plasticity of the TCR/pMHC trimolecular and pMHC bimolecular complexes by molecular dynamics simulations. The intermolecular contacts at the TCR/pMHC interface are similar for the complexes with the superagonist and the MBP self-peptide. The orientation angle between TCR and pMHC fluctuates less in the complex with the superagonist peptide. Thus, the higher structural stability of the TCR/pMHC tripartite complex with the superagonist peptide, rather than a major difference in binding mode with respect to the self-peptide, seems to be responsible for the stronger proliferative response.

## Introduction

Different antigen peptides presented by the same major histocompatibility complex (MHC; or in humans: human leukocyte antigen, HLA) protein can induce substantially different signals when recognized by the same T-cell receptor (TCR) (1). This observation is consistent with the role of hot spots at protein-protein interfaces as the antigen peptide is only a small part of the interface between TCR and pMHC. Furthermore, the antigen peptide represents a minor fraction of the tripartite complex which consists of more than 800 residues in the extracellular space (446 and 370 residues in the TCR and MHC protein studied here, respectively).

T-cell clone 3A6 was chosen as a biologically relevant and likely pathogenic autoreactive T-cell based on the following considerations. It has been isolated from a multiple sclerosis (MS) patient with relapsing-remitting MS, the most frequent form of the disease, and characterized in detail with respect to cytokine secretion (T helper 1* phenotype with secretion of interferon-gamma and less IL-17) (2, 3), HLA restriction by one of the MS-associated HLA-class II alleles (DR2a composed of DRA1*01:01 and DRB5*01:01) (2), and recognition of a wide range of variant peptides that result in agonist, partial agonist or even TCR antagonist responses (2). Furthermore, humanized mice expressing as transgenes the 3A6 TCR and the HLA-DR2a heterodimer develop spontaneous experimental autoimmune encephalomyelitis (EAE), the preferred animal model for MS (4). Finally, the trimolecular complex of the 3A6 TCR and DR2a/MBP-peptide allowed examining the structural interactions between the three molecules at 2.80 Å resolution and disclosed a very low avidity of the TCR interactions with the DR2a/MBP-peptide complex with no salt bridges and very limited hydrogen bonds at the contact points between TCR and MHC/peptide complex (5).

Here we investigate by means of microsecond molecular dynamics the structural stability (*i.e.*, kinetic stability of the bound state) and intrinsic flexibility of the tripartite complex 3A6-TCR/peptide/HLA-DR2a (abbreviated TCR/pMHC) and peptide/HLA-DR2a (pMHC). The choice of peptides was motivated by previously published surface plasmon resonance data which indicate that the HLA-DR2a MHC protein loaded with the superagonist peptide WFKLITTTKL has higher affinity for the 3A6-TCR and slower dissociation rate than the wild-type decapeptide MBP90-99 FFKNIVTPRT (called MBP-peptide or self-peptide in the following) (5). Furthermore, the superagonist shows two orders of magnitude higher proliferation of human CD4^+^T cell clone (called proliferative response in the following) than a single-point mutant of it (with Gly instead of Leu at the C-terminal residue, called peptide 28), and four orders of magnitude higher response than a three-point mutant of it (called peptide 36) and the wild-type MBP-peptide (6). These experimental data raise the following questions which inspired the simulations and their comparative analysis with the experimental observations. Is the footprint (intermolecular contacts) of the TCR on the antigen presenting surface of the pMHC complex different for peptides with different proliferative response? Does the tripartite complex with the superagonist peptide show significantly reduced plasticity than the complex with the self-peptide? Is the relative orientation of the TCR and pMHC restricted or does it fluctuate significantly, and how does it compare with the crystal structure?

MD simulations, despite known limitations such as accessible timescales and accuracy of the force fields, have provided significant insights in the flexibility of TCR/pMHC complexes (7–12). Recently, Fodor *et al.* (12) applied an ensemble enrichment method, which relies on multiple short MD simulations starting from X-ray diffraction data, to study conformational changes at TCR/pMHC interfaces. They extracted information of underlying dynamics overlooked in the comparison of crystal structures of TCR-bound and unbound pMHCs. Reboul *et al.* (13) obtained detailed information from MD simulations of two MHCs differing by a single polymorphism, with the same restricted peptide and the respective TCR/pMHC complexes. They observed transient contacts at the interface that are not present in the crystal structures. Furthermore, they related peptide fluctuations to the dynamic footprint of TCR/pMHC interactions, offering a pioneering model for TCR scanning of the pMHC surface. However, the role of flexibility and dynamics in TCR/pMHCs may remain system-specific and it is difficult to extract general binding mechanisms. As pointed out by Zhang and co-workers (14), there are few general rules also in describing the energetics of TCR/pMHC binding, similarly to the structural basis of TCR recognition. Other simulation studies of MHC class I and II dynamics have been reviewed recently (10, 15, 16). The novel aspect of our study is the comparative analysis of the TCR/pMHC complexes and pMHC complexes with four decapeptide antigens that show substantially different T cell signals despite their similar sequences. Our molecular dynamics simulations reveal a similar interface for the TCR/pMHC complexes with the different peptides, and a higher structural stability for the tripartite and pMHC complexes with the superagonist.

## Materials and Methods

The coordinates of the 3A6-TCR/MBP-peptide/HLA-DR2a complex were downloaded from the Protein Data Bank (PDB: 1ZGL (5)). The 1ZGL coordinate set lacks 19 residues which are localized in the MHC and TCR flexible loops and distant from the binding interface. The ModLoop server (17) (https://modbase.compbio.ucsf.edu/modloop) was employed to reconstruct the missing loops.

The HLA-DR2a restricted sequence of the wild-type MBP-peptide is FFKNIVTPRT (residues 90 to 99). The sequences of its mutants are shown in Figure 1. Simulations were carried out with capped peptides (acetyl and NH2 groups at the N-terminal and C-terminal residues, respectively) to emulate the peptides used in the T-cell proliferation assays (6). The side chain coordinates of the mutated residues were generated with the SwissPDBViewer software (18) and relaxed by means of the GROMACS software (19, 20). All simulated systems were generated from the 1ZGL structure, by removing the TCR and MHC coordinates for the free peptide runs, and the TCR coordinates for the pMHC runs. Hydrogen atoms were generated by the CHARMM-GUI (http://www.charmm-gui.org) server (21, 22). The simulations of the tripartite complex with the MHC loaded wild-type peptide were carried out with four different protonation states of the MHC (see Supporting Material). The analysis focuses on the state with all Asp side chains negatively charged while the remaining states are used mainly to evaluate the robustness with respect to the choice of protonation state.

**Fig. 1.**
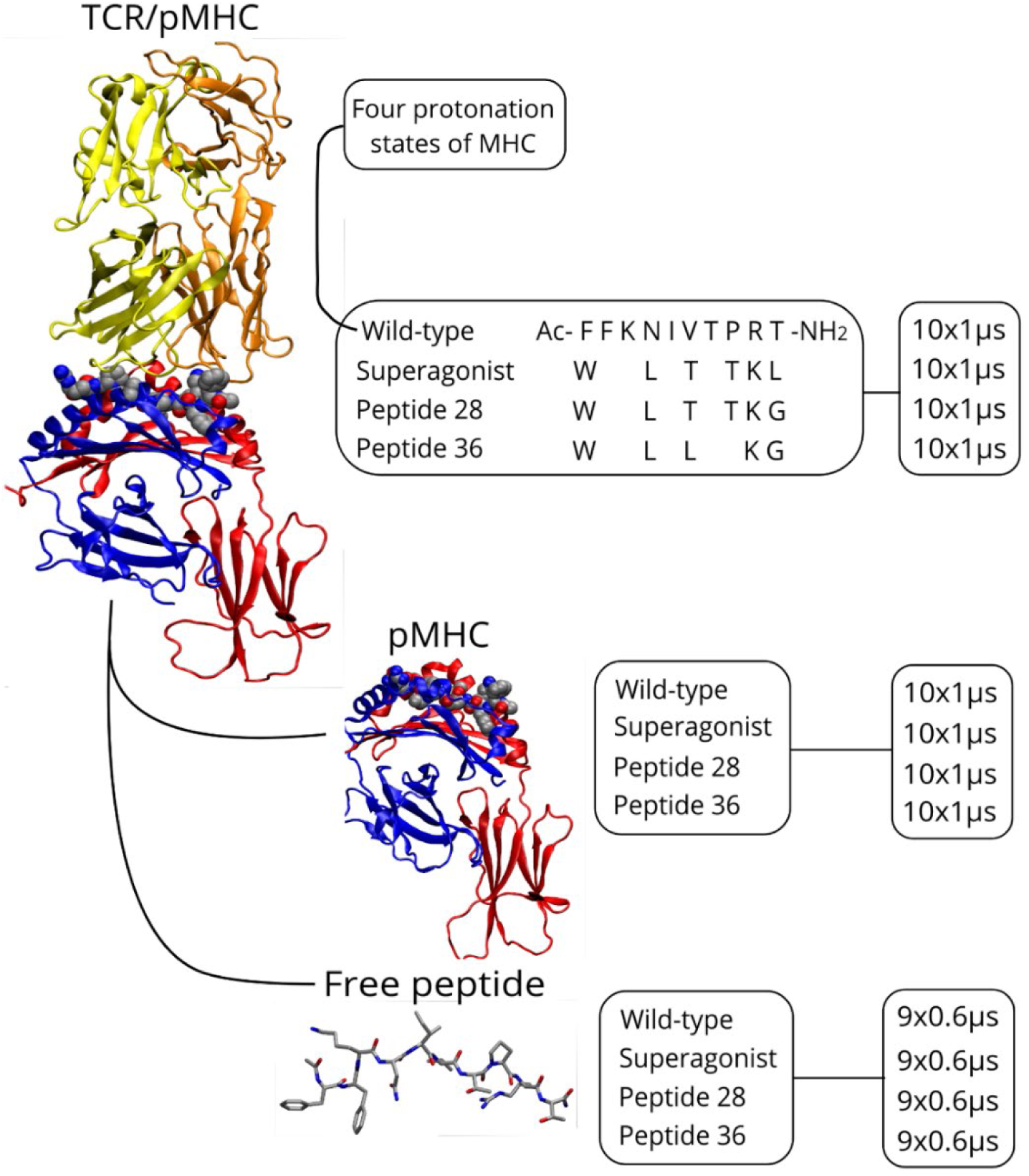
Overview of the simulations and sampling time. The PDB: 1ZGL (5) coordinate set was used to generate the starting structures for TCR/pMHC, pMHC, and free peptides. Complexes and free peptides were simulated in 10 and 9 independent copies, respectively. For the TCR/pMHC complex with the wild-type peptide, simulations were carried out with four different protonation states of the cluster of acidic sidechains in the groove of the MHC protein as a control to assess whether different protonation states result in higher variability in the wild-type with respect to the superagonist complex (see SI). For consistency with the T-cell proliferation assays, all peptides are decamers and have charge-neutralizing capping groups at the N-terminus (acetyl moiety, Ac) and C-terminus (NH2). The sequence of the MBP peptide and the mutated residues of the other peptides are shown. The effective concentration of the peptide at which 50% of the proliferative response is observed (EC50) is 34 ng/mL, 0.0034 ng/mL, 0.52 ng/mL, and 37 ng/mL for wild-type, superagonist, peptide 28, and peptide 36, respectively (6).

In all simulations, the CHARMM36 force field (23) was used with the TIP3P water model. Periodic boundary conditions were applied. The dimensions of the simulation boxes are 110×90×160 Å^3^for the tripartite complex, 83×85×110 Å^3^for the bipartite complex, and 60×60×60 Å^3^for the free peptides. The solvent contained K^+^and CL^−^ions, to have an ionic strength of 100 mM and an overall neutral system. The electrostatics forces were accounted for using the Particle Mesh Ewald (PME) algorithm. Truncation of all non-bonded interactions occurred at 12 Å and the LINCS algorithm was chosen to constrain all covalent bonds.

In every run, the energy minimization phase (steepest descent, convergence of maximum force under 100 kJ mol^−1^nm^−1^) was followed by two equilibration phases, each lasting 200 ps. In both phases, the temperature was held constant at 300K by an external bath with velocity rescaling and a coupling constant of 1 ps. In the first phase, the residues added in the loop reconstruction, the mutated residues and the solvent were relaxed in NVT ensemble while all remaining atoms of the solute were restrained. In the second phase, all sidechains were allowed to relax with the solvent, in NPT ensemble where the pressure was held constant at 1 atm by the Berendsen barostat, with a coupling constant of 2 ps. A 1-ns preproduction phase followed, in NPT ensemble (same settings for T and P as the previous phase), during which harmonic restraints (k=1000 kJ mol^−1^nm^−1^) were applied on two atoms of the TCR β chain, to prevent rotation of the solute and interactions of the complex periodic images. The atoms chosen were Cys92 and Cys148 Cα atoms, close to the central vertical axis crossing the system’s long dimension and localized in highly stable regions, *i.e.* two β-sandwiches characterizing each TCR chain. To set the volume for the NVT production runs, the average volume calculated from the second half of the 1 ns preproduction run was used, and the setting for the temperature was the same as before. The integration timestep was 2 fs and the saving frequency 50 ps, so that for each system a total of 200000 snapshots were saved along 10 independent 1 µs runs. All structural representations were produced using VMD 1.9.2 (24) and PyMol (PyMOL Molecular Graphics System, Version 2.7.2.1 Schrödinger, LLC).

## Results and Discussion

Experimental evidence has shown that the pMHC association is an obligate intermediate to the formation of the TCR/pMHC tripartite complex and eventual activation of immune response (25– 27). Thus, we decided to perform simulations of the free peptides, the pMHC complex, and the trimolecular complex. The summary of the simulations with total sampling time is depicted in Fig. 1. Simulations of the free TCR (3A6) and free MHC protein (HLA-DR2a) were not carried out as the main purpose of this study was to analyze the structural differences in the complexes that result in the variability of the proliferative response for the wild-type peptide and its mutants. Furthermore, experimental evidence indicates that HLA-DR2a is not stable in the absence of the peptide (28–30) and to date the peptide-unbound MHC (class I or II) has not been crystallized.

The results section starts with the analysis of the free peptides followed by the bimolecular and trimolecular complexes. Particular emphasis is given to the comparison between the wild-type decapeptide MBP90-99 (FFKNIVTPRT) and superagonist peptide (WFKLITTTKL), as the latter shows a four orders of magnitude higher proliferative response than the former (6). Of note, surface plasmon resonance measurements have provided direct evidence of the higher affinity and slower dissociation rate from the TCR for the MHC loaded with the superagonist than the wild-type peptide (5).

### Simulations of the unbound peptides

Simulations with the individual peptides in solution were carried out to analyze potential differences in their preferred conformations. Eventual differences would influence the formation of the pMHC complex directly but are expected to have a marginal influence on the tripartite complex. Nine independent 0.6-microsecond runs were carried out for each of the four 10-residue peptides (Fig. 1, bottom). The peptides populate very similar regions of conformational space in their unbound state. The distributions of radius of gyration (Rg) and end-to-end distance show that they sample extended conformations, with end-to-end distances peaked in the range of 22 Å to 28 Å (Fig. S1). The two-dimensional histograms of normalized radius of gyration (measure of size) versus asphericity (measure of shape) provide further evidence that the free peptides are most of the time in an extended state but can also populate compact conformations (Fig. 2). The peptides are almost fully extended in the pMHC bipartite complex (purple in Fig. 2A) and TCR/pMHC tripartite complexes (blue in Fig. 2B). A direct comparison of the wild-type peptide and the superagonist indicates that the bound state of the latter overlaps with the extended portion of the free state, while the wild-type peptide shows a somewhat larger asphericity in the bound than the free state (top panels in Fig. 2A and 2B). Thus, according to these geometric variables the superagonist peptide is subject to a lower degree of structural rearrangement in the formation of the bipartite and tripartite complexes. Note that only the peptide backbone atoms were used in the calculation of geometrical properties to have a common measure of peptide conformations not influenced by differences in side chains. To analyze the statistical uncertainty associated with the free peptide simulations, we plot the 2D-histograms of each independent run. The similar 2D-histograms give evidence of statistical robustness in the sampling of the extended state in all peptides (Fig. S2). The simulation results of the free peptides cannot be compared with cellular assays or *in vivo* results as the mechanism of peptide loading on the MHC is a complex scenario, in which peptide exchange catalyzed by HLA-DM and conformational plasticity of the MHC molecule play essential roles (10, 31–34).

**Fig. 2.**
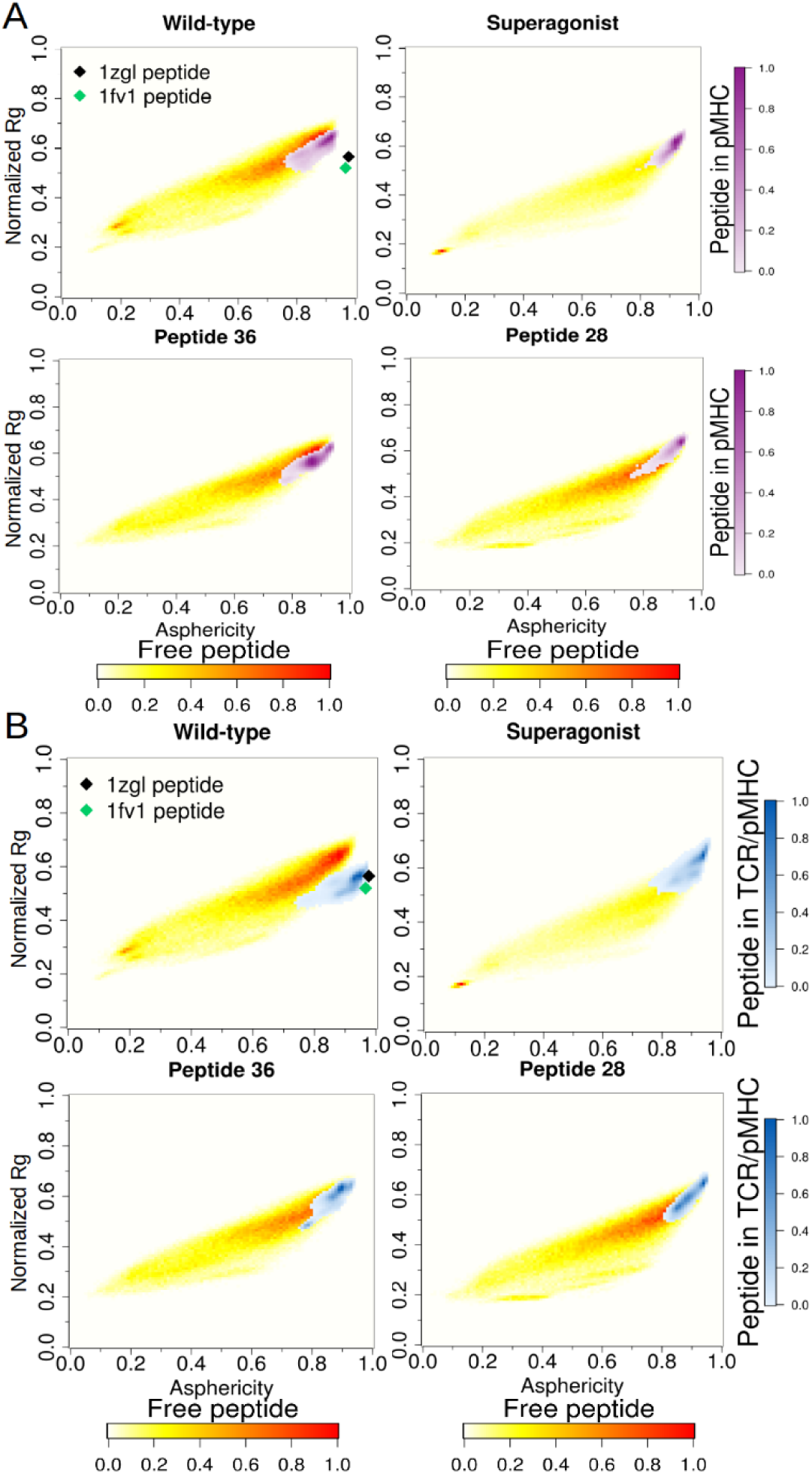
Two-dimensional histograms of normalized radius of gyration (Rg) and asphericity. Linear color scales refer to the free peptides (yellow to red in panels A and B), the peptides in the simulations with the bipartite pMHC complex (purple in A), and in the simulations with the TCR/pMHC complex (blue in B). The conformations of the MBP decapeptide in the TCR/pMHC and pMHC crystal structures (black and green diamonds, PDB: 1ZGL (5) and 1FV1 (55), respectively) are close to those sampled in the simulations, with slightly higher values of asphericity.

### Simulations of the pMHC bimolecular complex

#### Structural stability

Ten independent 1-microsecond runs were carried out for each of the four pMHC complexes (Fig. 1). As mentioned above, this subsection focuses on the simulations of the pMHC bipartite system while the comparative analysis with the TCR/pMHC trimolecular system is presented in the next subsection. The temporal series of the root mean square deviation (RMSD) of the Cα atoms from the equilibrated structures show that, irrespective of the peptide sequence, the pMHC structure is stable over the 1-microsecond time scale of the individual simulations (Fig. 3A). The mean values and standard error of the RMSD along the total sampling of 10 microseconds are 2.3±0.2 Å, 2.4±0.2 Å, 2.5±0.4 Å, and 2.7±0.4 Å, for the pMHC complexes with superagonist, peptide 28, peptide 36, and wild type, respectively (here and in the following text the standard error for each system is evaluated as the standard deviation of the ten average values along each independent run). The higher structural stability of the pMHC complexes with superagonist compared to wild-type is consistent with the stronger proliferative effect of the superagonist, as the rigidity of the pMHC complex is expected to contribute to the enthalpic stabilization of the TCR/pMHC assembly. On the other hand, a ranking of the four peptides is not possible as the differences are small and within the statistical uncertainty. While the structural rigidity of the pMHC would result in a higher entropic cost of association to the TCR, several studies of TCR/pMHC complexes have reported on the importance of enthalpic contributions (35–37). The thermodynamic stability of the complex is the result of both entropic and enthalpic factors. It remains difficult to identify thermodynamic signatures in the TCR/pMHC complex formation (38, 39), as its trimolecular nature translates into a wide range of entropic cost *versus* intermolecular contacts contributions. This has outcomes in TCR degeneracy and cross-reactivity (40–42).

**Fig. 3.**
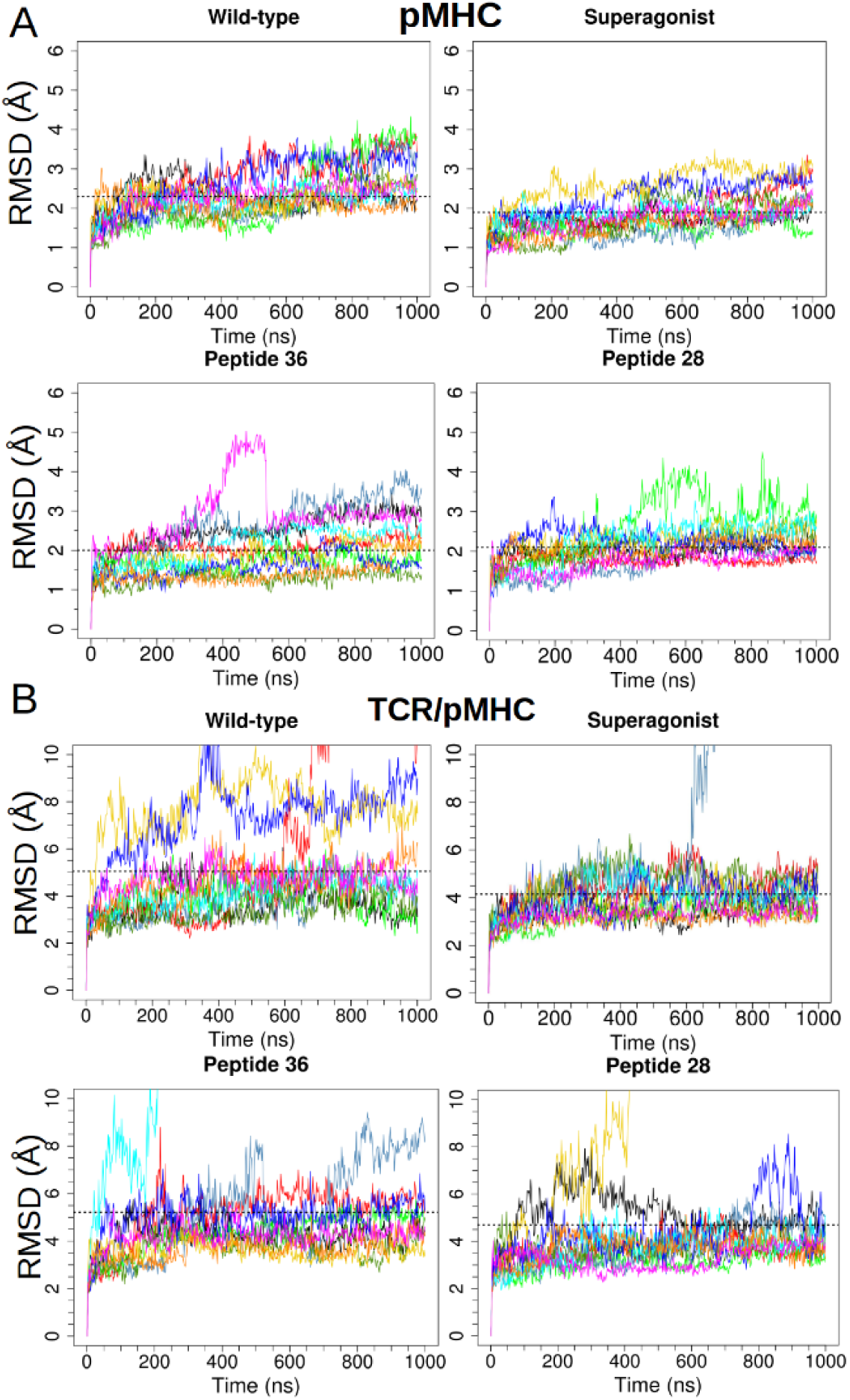
Structural stability of the pMHC complex (A) and TCR/pMHC complex (B). The time series of Cα RMSD from the equilibrated structure are shown with different colors for the individual runs. The average over all of the sampling for each system is shown (dashed line).

The two long α-helices of the MHC protein are located in the domains α1 and β1; they surround the peptide binding groove and contact both peptide residues and CDR (complementarity-determining region) loops of the TCR. To inspect the plasticity of the binding site on the MHC proteins, we have analyzed the time series of RMSD for the recognition helices, after initial fitting to the MHC α1β1 domains including the two α-helices (Fig. S3). The RMSD values of the recognition helices are very similar for the four complexes. Again, a slightly reduced flexibility is observed for the superagonist peptide, which has the lowest average of the RMSD at 1.9±0.3 Å, while peptide 28, wild-type and peptide 36 have averages of 2.1±0.3 Å, 2.3±0.2 Å, and 2.0±0.5 Å, respectively.

Taken together, the simulation results suggest that the superagonist peptide stabilizes a pMHC surface optimized for contacts with the TCR. It is known that motions of both peptide and MHC influence binding of TCR and could have a role in “tuning” the T cell response (43–45). Insaidoo *et al.* (46) have used structural data, MD simulations and dissociation kinetics to show how antigen peptide modifications affect pMHC flexibility and consequently TCR binding. In their study, higher pMHC flexibility induced a weaker TCR binding, while maintaining a strong binding of the modified antigen to the MHC molecule.

#### Peptide/MHC interface

It is useful to analyze intermolecular contacts at the binding interface as they contribute most to binding of the peptide. The pattern of peptide-MHC contacts is plotted as the frequency of each contact averaged over 10 simulations and displayed as heat map with linear color scale (Fig. 4A). The four peptides show very similar patterns of contacts with the MHC protein (Fig. S4). The two major stripes of contacts reflect the parallel arrangement of the peptide backbone with the MHCα1 helix and the antiparallel arrangement with the β1 helix, respectively. The binding pockets for anchor peptide residues are well documented for the MHC chains of the HLA-DR2a haplotype (47–49). In detail, the binding motif on the MHC class II protein consists of pocket I, which accommodates large aromatic residues (conserved Phe2 in the simulated peptides); pocket II, that binds aliphatic residues (conserved Ile5); and pocket III, which is formed by a cluster of acidic side chains and binds a conserved basic residue in position 9, *viz*., Arg9 in wild-type and Lys9 in the three mutated peptides (pocket numbering after reference (47)). It emerges from the contact maps that the N-terminal acetyl moiety establishes hydrophobic interactions with MHCα1 and β1 regions, *i.e.*, in the same hydrophobic pocket that accommodates the N-terminal residue. This cannot be compared directly with the crystal structure as the peptide is capped by charge-neutralizing groups in the simulations, while it was uncapped in the crystallization experiments. Interestingly the proliferation assay with Ac-decapeptide libraries (6) provides evidence that the acetylation enhances peptide binding to MHC and thus is likely to increase T-cell activation.

**Fig. 4.**
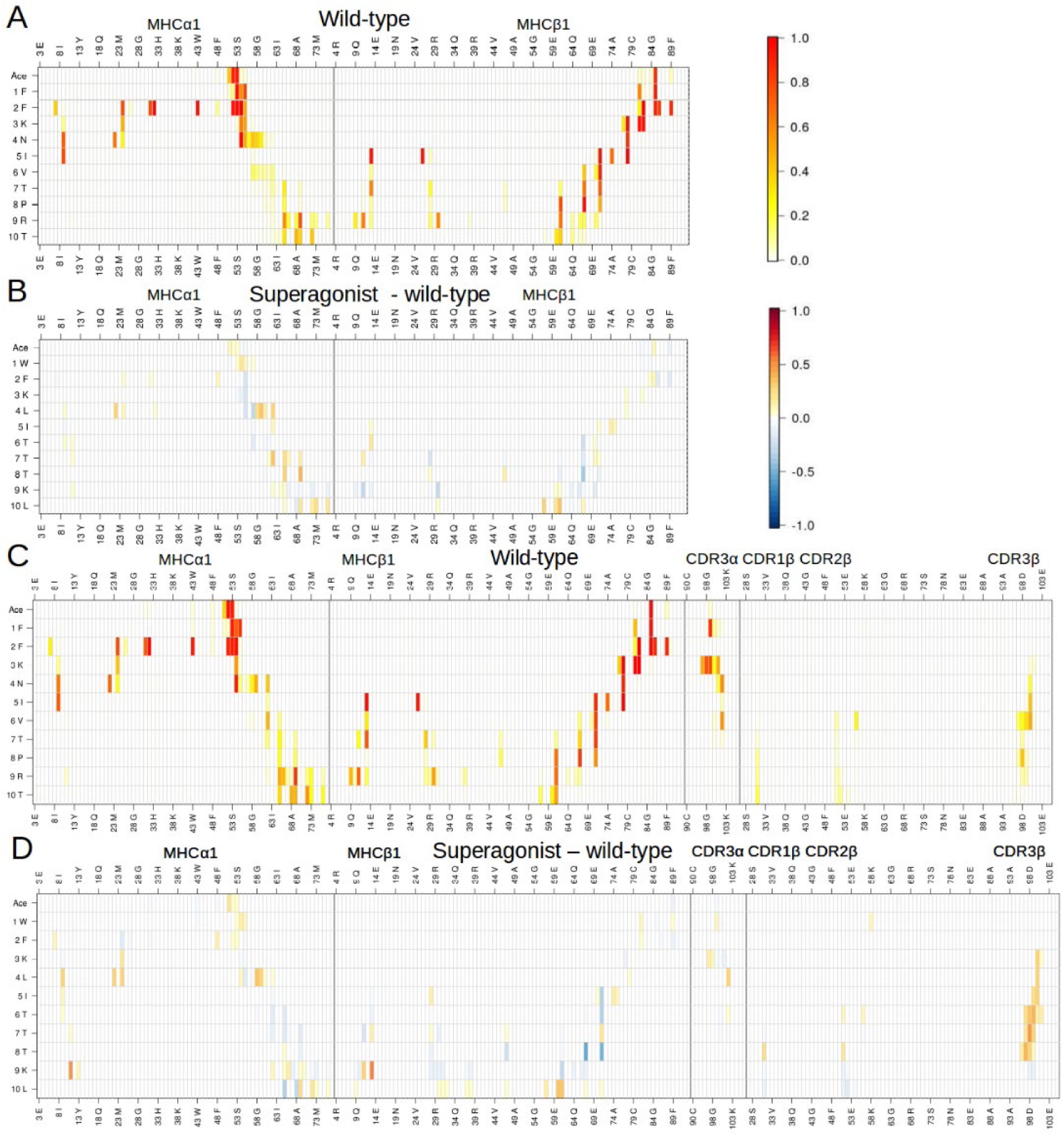
Contact maps for the pMHC runs (A,B) and TCR/pMHC runs (C,D). In all plots, peptide residues are listed on the vertical axis, while the interacting MHC and CDR loops residues are on the horizontal axis, separated by vertical lines. Here, a contact represents pair of atoms within a 3.5 Å distance. (A,C) The frequency of contact is normalized from 0 to 1 and represented as a heat map, with linear color scale. (B,D) The difference map is calculated by subtracting the contact map of the complex with the MHC loaded with wild-type from the complex with the superagonist. Thus red represents a higher frequency of contact for the MHC loaded with superagonist than wild-type peptide.

For a direct comparison between different peptides it is useful to calculate maps of differences in individual contacts. The map of contact differences between bipartite superagonist and wild-type (Fig. 4B) shows a stronger binding to the MHCα1 domain of the superagonist central residues, specifically Leu4, Thr7 and Thr8. Furthermore, Leu10 in the superagonist forms more contacts with the MHC protein than Thr10 of wild-type, indicating that the hydrophobic interactions with both MHCα1 and β1 allow to stabilize the superagonist C-terminal end and contribute to the stability of this complex. While the superagonist and the wild-type peptide differ in six residues, there is only a single-point difference between superagonist and peptide 28 (Leu10Gly). Their map of contact differences illustrates that the single mutation at the C-terminal residue influences not only the contacts between the last residue and the MHC protein, but also those of preceding residues that are identical in the two peptides (Fig. S5a). Thus, a single difference between two peptides can have effects that propagate to contacts involving residues distant in sequence and space from the mutated one. A large-scale study with extensive sampling by Ayres *et al.* (11) has analyzed the fluctuations of nonameric peptides restricted by MHC class I molecules to generate a model for peptide flexibility based on peptide sequence and chemical composition. The relationship between peptide sequence and propagation of fluctuations, as observed also in our study, is of interest when considering antigen immunogenicity.

### Simulations of the TCR/pMHC trimolecular complex

#### Structural stability

Ten independent 1-microsecond runs were carried out for each of the four tripartite complexes (Fig. 1). The time series of RMSD from the equilibrated structures of the TCR/pMHC complex show similar flexibility irrespective of the bound peptide (Fig. 3B). The values of RMSD oscillate mainly between 3 Å and 5 Å except for one run of each system (two runs for the complex with the wild-type peptide) in which the RMSD reaches values larger than 10 Å (Fig. S6). The mean values and standard error of the RMSD along the total sampling of 10 microseconds are 4.2±1.1 Å, 4.7±2.6 Å, 5.1±1.6 Å, and 5.2±2.6 Å, for the TCR/pMHC complexes with superagonist, peptide 28, wild-type and peptide 36, respectively. The statistical error does not allow one to rank the four peptides. Nevertheless, the lowest mean value for the superagonist, *i.e.*, highest structural stability, is not inconsistent with its strongest proliferative effect. It is important to note that the structural stability of the complex with the superagonist is not an artifact due to the modeling of the six side chains that differ between this peptide and the wild-type because the superagonist was modeled starting from the crystal structure of the wild-type and manual building of side chains usually results in larger deviations during the molecular dynamics runs.

The relative displacement of the TCR with respect to the pMHC complex can be visualized upon structural overlap with the backbone of the latter (Fig. 5A,B). To quantitatively monitor this displacement, we plot the time series of RMSD of the TCR variable regions (VαVβ) after initial alignment on the peptide and MHCα1β1 domains. They show that the motion of the TCR is lowest for the tripartite complex with the superagonist (Fig. 5C). The mean value and standard error of the RMSD along the total sampling of 10 microseconds are 5.4±1.3 Å, 8.3±8.7 Å, 7.6±3.3 Å, and 5.6±1.5 Å, for the pMHC complexes with superagonist, peptide 28, wild type, and peptide 36, respectively. There are three runs of the wild-type that deviate most. They are characterized by the loss of contacts between the TCR, either α or β chain, and the contacting regions on the MHC, *i.e.* β1 and α1 helix respectively (Fig. S6).

**Fig. 5.**
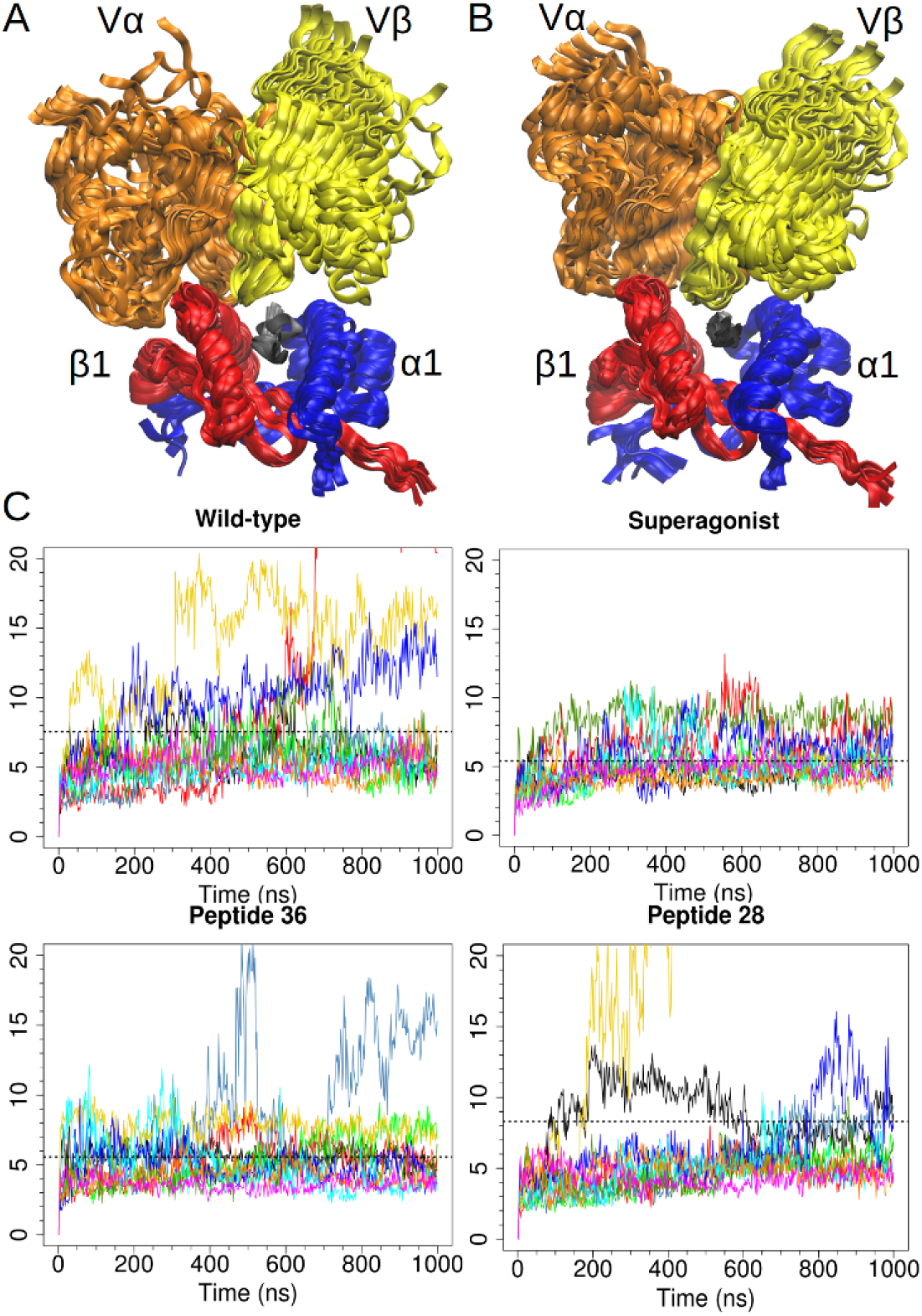
Displacement of TCR. (A) Overlap of 11 molecular dynamics snapshots of wild-type complex aligned on the peptide and MHCα1β1 domains (Cα atoms), to visualize the flexibility of the TCR VαVβ chains. The snapshots were taken from a 1 μs run at equally spaced intervals of 100 ns starting from the initial structure. (B) Same as A for the complex with the superagonist. The complex with the wild-type peptide shows a slightly higher flexibility, compared to the superagonist complex. (C) Cα RMSD time series for TCR VαVβ after initial alignment on the pMHCα1β1. The average over all of the sampling for each system is shown (dashed line). The superagonist complex shows the smallest RMSD values.

#### TCR/pMHC orientation angle

To further quantify the TCR displacement with respect to the pMHC complex we plot the time series and distribution of the TCR to pMHC orientation angle which is defined as the angle between the vector connecting the termini of the peptide and the vector connecting the centers of mass of the TCR Vα and Vβ domains (Fig. 6A) (50, 51). The orientation angle shows substantially higher fluctuations for the complex with the wild-type peptide than with the superagonist (Fig. 6B,C; see also Fig. S15 for the comparison with the wild-type peptide in the complex with the MHC in four different protonation states). Over the whole sampling of 10 microseconds, the average value and standard error are 72.4±7.8° and 75.5±3.4° for the wild-type and superagonist, respectively (the standard deviation over the total sampling of 10 microseconds is 11.0° and 5.7°, respectively). Furthermore, there are two runs of the wild-type in which the orientation angle reaches values smaller than 40°, during which the TCR tends to a parallel orientation with respect to the peptide.

**Fig. 6.**
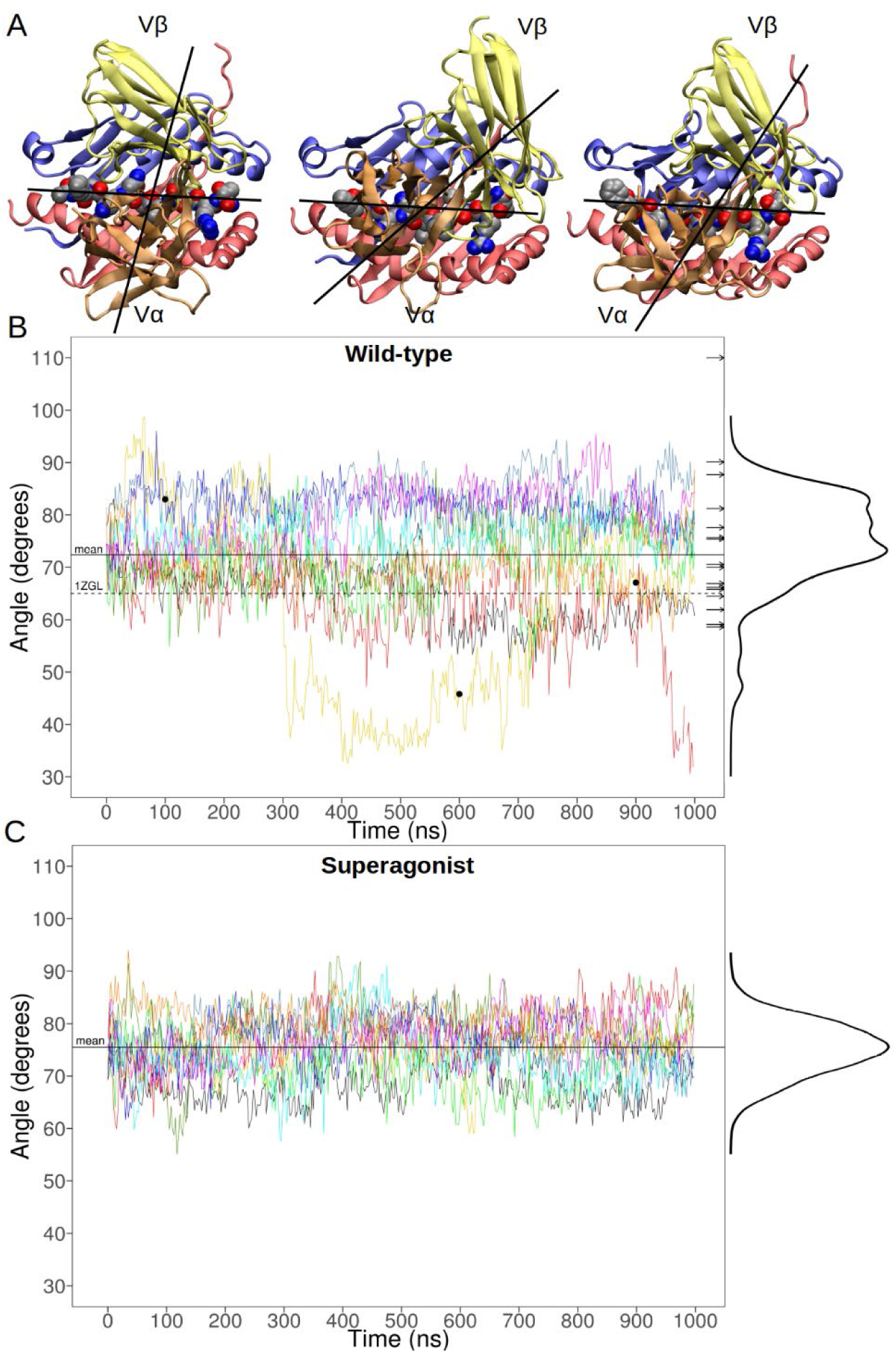
Orientation angle between TCR and pMHC surface. (A) The orientation angle is defined by two vectors: one is traced through the centers of mass of TCR VαVβ domains and the second connects the Cα atoms of terminal residues 1 and 10 of the peptide. The three representations are snapshots from a wild-type run with large variations in the angle. (B,C) Time series of individual molecular dynamics runs (colored lines) and the histogram (right margin). The wild-type complex (B) shows a broader range of rotational freedom than the superagonist (C).The black circles in B indicate the position along the time series of the snapshots shown in A. The orientation angles measured in different crystal structures (represented by arrows) fall within the range sampled by the wild-type simulations (B), irrespective of the MHC class. The PDB IDs of the crystals used are 1BD2, 1MI5, 3RGV, 4JFF, 4JRX, 4MNQ, 4PRI, 5BRZ, 5HHO for MHC class I, and 1FYT, 1J8H, 1YMM, 2IAN, 3TOE, 4C56, 4GRL for MHC class II. Only the Ob.1A12/MBP/DR2b (1YMM) complex is distant from the sampled range, with a 110° orientation angle.

The comparison with the TCR to pMHC orientation angle as measured on a set of 16 crystal structures (containing nine and seven MHC proteins of class I and II, respectively) reveals that the range of values observed in different crystals is sampled along the simulations (Fig. 6B). The only exception is the value of the orientation angle in the Ob.1A12/MBP/DR2b crystal structure (PDB: 1YMM, associated with autoimmune response) which has an orientation angle of 110° outside the range of both MHC classes, and exhibits a highly asymmetrical TCR/pMHC interaction (52). The broad range of orientation angle values sampled along the simulations indicates that care has to be taken in comparing individual values of the angle as measured on crystal structures, which do not capture the rotational motion of the tripartite complex.

#### TCR/pMHC interface

The pattern of contacts between the TCR and pMHC is very similar in the four complexes and for all of the three interfaces, *viz.*, peptide with MHC, peptide with TCR (Fig. 4C,D and S7, S8), and MHC with TCR (Fig. 7 and S9). It is interesting to note that there are only minor differences in the contacts between peptide and MHC in the presence or absence of the TCR (compare Fig.4A and 4C) which indicates that the binding of the latter does not modify the interface between peptide and MHC.

**Fig. 7.**
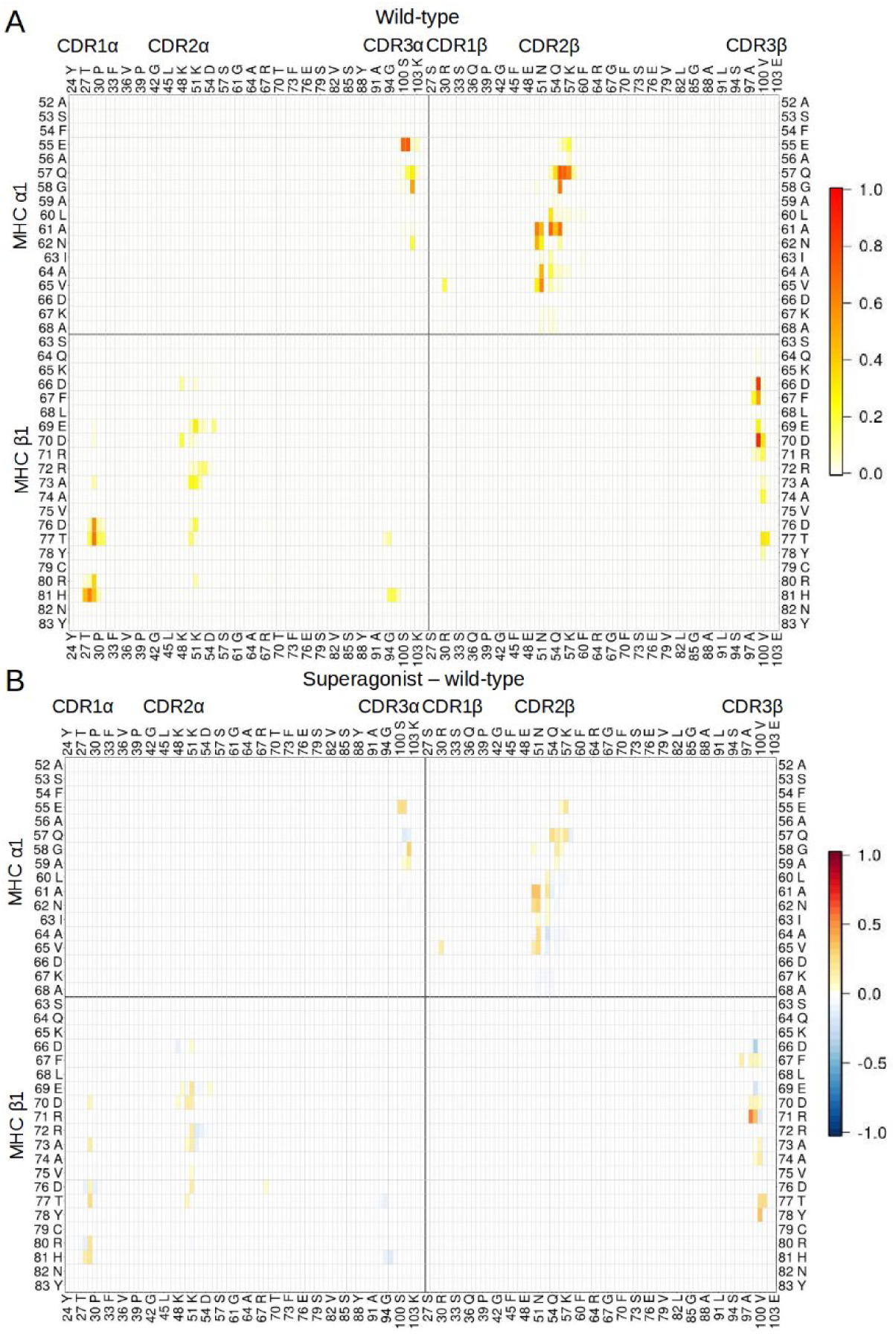
Contact maps for the TCR/pMHC interface. (A) Frequency of contacts in the complex with the wild-type peptide, averaged over 10 simulations, displayed as heat map with linear color scale. The contact interface between the two proteins is limited and a similar pattern is found for the other peptides. (B) Differences at the TCR/pMHC interface between superagonist and wild-type complexes, with linear color scale, where red indicates stronger binding for the tripartite complex with the superagonist and blue for the wild-type. The superagonist stabilizes the interface more than the wild-type peptide, in particular the contacts between CDR3β loop and MHC β1 helix.

The superagonist peptide shows stronger contacts with the TCR than the wild-type, particularly with the CDR3β loop (Fig. 4D). The stronger interactions involve mostly the central segment of the peptide (residues 3-8 with CDR3β), while limited differences are found in the contacts at the C-terminal segment, *i.e.* Lys9 and Leu10, with few residues on both MHC chains. On the other hand, the stretch of three threonines of the superagonist shows less interactions with the MHC than the wild-type. It is interesting to compare the superagonist and peptide 28 which differ only at the C-terminal residue. The superagonist establishes more persistent contacts with its Leu10 and Lys9 and the MHCα1β1 domains when compared to peptide 28 (Fig. S8c). In addition, there are more interactions with the CDR3β for most of the residues. These simulation results confirm that a single-point mutant (Gly10Leu) can have a stabilizing effect on the binding interface that is propagated to other segments of the peptide, as already observed in the analysis of the pMHC bipartite complex. Previous studies by others have discussed how small residue changes at TCR/pMHC interfaces can lead to differences in affinity and signal propagation, while maintaining an overall conserved binding mode (13, 53, 54).

Except for very few contacts spread over the interface of the two proteins, the contacts between MHC and TCR are in general more stable in the tripartite complex with the superagonist than the wild-type peptide (Fig. 7B). In particular, the complex with the superagonist shows stronger CDR3β–MHCβ1 helix interaction than the complexes with wild-type peptide or peptide 28 (Fig. 7B and S9). Between superagonist and wild-type the difference is mainly in the stronger Arg71(MHCβ)– Asp98(CDR3β) contact. The above observations suggest that the distinctive feature of the superagonist peptide is to induce more contacts of the CDR3β loop on both the peptide central region and MHC helices. The statistical robustness of the contact maps is supported by a block averaging analysis, which shows for each complex very similar results between two randomly chosen subsets of five trajectories (Fig. S10 and Fig. S11).

#### TCR imprint on the pMHC

The contact maps illustrate the time-averaged values over the whole sampling for individual pairs of residues. Complementary information is contained in the time series of the surface buried at the interface which shows the temporal evolution of the TCR imprint on the pMHC. The buried surface area (BSA) was calculated along the TCR/pMHC runs. Overall, average values of BSA and fluctuations are similar for the four peptides (Fig. S13). A higher buried surface is observed for the than the wild-type peptide, with average values of 1054±134 Å^2^and 932±131 Å^2^, respectively. The superagonist adapts the binding interface to increase the imprint of the TCR on the pMHC. This results in a larger enthalpic contribution to the association of the receptor. Of note, the BSA calculated for the crystal structure (1020 Å^2^) is within the fluctuations observed along the simulations for the four complexes.

#### Widening of MHC binding groove upon TCR association

The crystal structure of the bipartite complex MBP/HLA-DR2a (PBD: 1FV1 (55)) can be compared with the 3A6/MBP/HLA-DR2a structure to identify rearrangements in the pMHC surface upon TCR engagement (5). A widening of the HLA-DR2a binding groove is observed in the crystal structure of TCR 3A6-bound state. The MHCβ1 helix shows the largest displacement at Asp66β (Fig. 8A). The distributions of distances between the Cα atoms of Asp66β and Val65α for all bipartite and tripartite runs show a smaller aperture of the MHC binding groove in the pMHC than the TCR/pMHC complexes in agreement with the respective X-ray structures (Fig. 8B). To monitor the widening of the MHC groove we also show the temporal series of the distance in the wild-type and superagonist peptide (Fig. 8C,D). The latter shows a persistently greater aperture of the MHC binding groove in the TCR-bound state than the wild-type. This result, together with the slightly higher stability of the pMHC (Fig. 3A) and the recognition helices (Fig. S3) in the superagonist runs, may be indicative of the “tuning” of TCR recognition by the pMHC conformational dynamics. Hawse and co-workers (56) have discussed how TCRs may scan for ligands and how they could be facilitated in binding to pMHCs that best match the structural and motional properties of the receptor. They observe that the sampled conformations and the rates at which the peptide and CDR3β move in the free TCR and in the free pMHC are similar, and this promotes trimolecular complex formation.

**Fig. 8.**
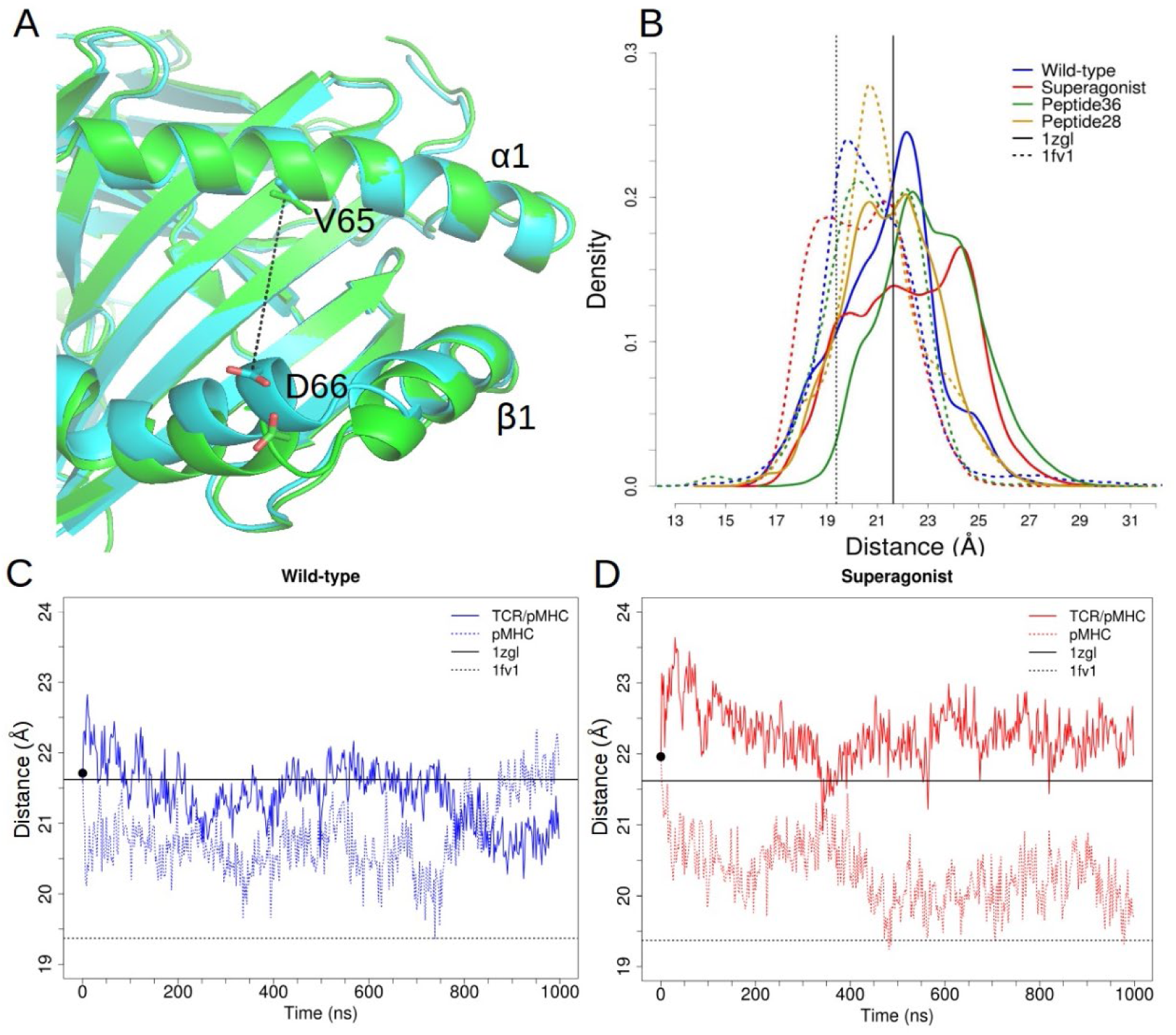
Aperture of the MHC binding groove. (A) The MHC structures from the pMHC complex (cyan, from PDB 1FV1) and TCR/pMHC tripartite complex (green, from PDB 1ZGL) are shown in the overlap based on Cα atoms. The distance between the Cα atoms of Asp66β and Val65α (black dashed line) is used to monitor the aperture. (B) Distribution of the Asp66β-Val65α distance for the runs with TCR/pMHC (solid line) and pMHC (dashed lines). The distances measured in the 1ZGL and 1FV1 crystal structures are shown as a basis of comparison (solid and dashed vertical lines at 21.6 Å and 19.4 Å, respectively). The widening of the MHC groove in the different TCR/pMHC structures is evident, especially for the superagonist complex. (C,D) Time series of the Asp66β-Val65α distance averaged over 10 simulations, for wild-type complex (C) and superagonist complex (D). The tripartite and bipartite runs (solid and dashed lines, respectively) have a common starting distance (black point) as they were generated from the same crystal. The runs with the bipartite pMHC complex evolve to a binding groove with smaller aperture close to that observed in the pMHC crystal structure 1FV1.

To assess the statistical significance of the data on the MHC aperture and the robustness with respect to the choice of protonation state of the MHC protein we have compared the distribution of the Asp66β-Val65α distance for the four different protonation states of the MHC protein in the complex with the wild-type peptide (Fig. S12). The distributions show that irrespective of the protonation state the binding groove is larger in the tripartite system than the pMHC bimolecular complex.

## Conclusions

We have used explicit solvent molecular dynamics simulations to analyze the influence of the antigen peptide on the structural stability of the TCR/pMHC (class II) complex, 3A6-TCR/MBP-peptide/HLA-DR2a. Ten 1-μs molecular dynamics runs were carried out for the TCR/pMHC and pMHC complexes with the wild-type peptide (Ac-FFKNIVTPRT-NH2, *i.e.* residues 90-99 in the MBP sequence) and with three peptide mutants. The pairwise differences for the four peptides are of 1, 2, 3, 5, 6, and 6 residues, respectively (Fig. 1). Previous *in vitro* studies have reported slower dissociation rates for the tripartite complex with the superagonist peptide than the wild-type peptide (5). Furthermore, significant differences in the proliferative response have been measured with up to four orders of magnitude stronger response for the superagonist (Ac-WFKLITTTKL-NH2) with respect to the wild-type peptide (6). A surprising experimental observation is that peptide 28, which differs from the superagonist only by the single point mutation Leu10Gly, shows a two orders of magnitude weaker response than the superagonist, whereas peptide 36 (Ac-WFKLILTPKG-NH2) and the wild-type peptide showed similar response despite half of their residues differing. Unfortunately, surface plasmon resonance data have not been reported for peptides 28 and 36.

Four main observations emerge from the analysis of the molecular dynamics trajectories. (*I*) The simulations of the wild-type peptide and three mutants in the unbound state show similar free-energy projections onto geometric variables that monitor compactness and deviation form spherical shape. All peptides populate the extended state which is the one observed in the pMHC complexes, and the superagonist has a slightly larger overlap of free and bound states than the wild-type peptide. (*II*) The contact maps along the simulations are very similar irrespective of the peptide sequence, except for a stabilization of the interactions between peptide and TCR CDR3β loop for the superagonist complex. Even a single residue difference (Gly10Leu) at the C-terminus of the peptide results in stronger peptide/TCR CDR3β contacts for the superagonist with respect to peptide 28. (*III*) The four tripartite complexes, which differ only in the antigen peptide, have similar flexibility with a more pronounced structural stability for the superagonist than the wild-type peptide. This difference is consistent with the slower dissociation rate of the TCR from the superagonist/HLA-DR2a complex than from the wild-type peptide/HLA-DR2a complex as measured by surface plasmon resonance (5). (*IV*) The tripartite complex with the superagonist shows the smallest fluctuations in the orientation angle of TCR to pMHC. It is interesting to note that on a microsecond time scale the fluctuations of the orientation angle cover the range of values of the available TCR/pMHC X-ray structures. This simulation result indicates that individual crystal structures have to be considered as snapshots of an intrinsically flexible system which is frozen in a crystalline arrangement (12, 13). Thus, care has to be taken in using only crystal structures for drawing conclusions on the influence of the orientation angle and/or other structural features (*e.g.*, surface buried at the interface) on T-cell signal propagation.

There are two main limitations in the present study. First, the individual simulations of the complexes reach a time scale of 1 µs (with cumulative sampling of 10 µs for each system), while there is the possibility that major rearrangements take place on longer time scales. However, the essentially identical crystal structures of four TCR/pMHC class I complexes that mediate very different T cell responses (53) provide evidence that the simulation results are valid even at longer time scales.

The second limitation concerns the truncated system investigated here. The present simulation study started from the crystal structure of the tripartite complex which includes only the α and β ectodomains of the TCR and did not take into account the other domains and the transmembrane segments. As such, it is not possible to analyze the broader range of structural rearrangements of these proteins (influenced also by co-receptors) that are linked to T-cell response (57–60). However, the simulation results for the wild-type peptide and superagonist are consistent with the thermodynamics and kinetics of TCR/pMHC dissociation as measured by surface plasmon resonance and the large difference in proliferation effects.

In conclusion, our simulation study suggests that the high proliferative response of the superagonist, which differs from the MBP self-peptide mainly in the residues that are in contact with the 3A6 TCR, is due to higher rigidity of the TCR/pMHC complex rather than substantial differences in binding mode. It remains to be investigated if the observed similarities and differences are valid for other superagonist peptides.

## Author contributions

A.C. and R.M. designed the study. I.S. carried out the simulations. I.S. and A.C. analyzed the simulations.

