## Supporting Material for "The TCR/peptide/MHC complex with a superagonist peptide shows similar interface and reduced flexibility compared to the complex with a self-peptide"

### Supporting Methods

#### Simulations of different MHC protonation states

The simulations of protonated systems, *i.e.* addition of a single proton to individual acidic residues to neutralize their charge, were realized as a control to measure the structural flexibility of the wild-type complex in other possibly existing states. In our system, relevant protonation involves the residues of the MHC acidic cluster, which are key to binding the basic aminoacid in position 9 of wild-type (*i.e.* Arg9) and mutated peptides (*i.e.* Lys9). The cluster involves up to 6 residues: one glutamate and five aspartates. The protonation of these closely spaced residues has consequences on their p$K_a$ values and thus on the interaction with the peptide residue 9. The residues were chosen after analyzing the p$K_a$ through PROPKA calculations on the PDB2PQR server (Dolinsky, T.J., J.E. Nielsen, J.A. McCammon, and N.A. Baker. 2004. PDB2PQR: an automated pipeline for the setup of Poisson-Boltzmann electrostatics calculations. Nucleic Acids Res. 32: W665–W667.). This method allows rapid, empirical predictions of p$K_a$ values taking into account the protein structure. Specifically, the p$K_a$ calculations were repeated for the wild-type crystal and different equilibrated structures. The parameters chosen were the CHARMM force field and pH 7, used to assign the p$K_a$ values. From this analysis we identified residues of the MHC acidic cluster with large shifts in p$K_a$ with respect to the standard value of their species, *i.e.* 3.8 for aspartate and 4.5 for glutamate. The initial values of p$K_a$, when the residues were chosen for protonation, are listed in Table S1; the values will change during the simulations due to movement of residues and their non-bonded interactions. We have selected for mutually exclusive protonation the residues Asp66 (MHCα), Asp11 (MHCβ), and Asp37 (MHCβ), which exhibited the highest shifts of p$K_a$ in the X-ray and energy minimized structures. Also, visual inspection evidences the proximity of Asp66α and Asp11β to Arg9 in the peptide. We thus obtained a set of four protonation states (including the original unprotonated system) for the TCR/pMHC with the MBP-peptide. The addition of the proton was realized through the CHARMM-GUI server (22), specifically using the PDB manipulator function (23), to generate the starting structure for the GROMACS simulations. The sampling time of each system is listed in Table S2.

The RMSD analysis on the set of protonation complexes shows that they are subject to a similar degree of variability as the original wild-type complex (Fig. S14 in the Supporting Material). Specifically, the average values and standard errors of RMSD of Cα for the whole system (Fig. S14a) are 5.1±1.6 Å, 5.2±2.9 Å, 4.0±1.0 Å, and 5.0±1.8 Å, for wild-type, Aspp11, Aspp66 and Aspp37, respectively. The results of the RMSD for TCR VαVβ atoms after fitting to MHCα1β1 (Fig. S14b) are 7.6±3.3 Å, 7.6±3.2 Å, 6.2±2.8 Å, and 8.1±5.7 Å, for wild-type, Aspp11, Aspp66 and Aspp37, respectively. Thus, when comparing the four control simulations and the superagonist complex, the latter shows lower flexibility. The same comparison was made with other measures we discuss throughout the text, relevant for the analysis of the TRC/pMHC: the orientation angle (Fig.S15 in the Supporting Material) and the buried surface area (Fig.S16 in the Supporting Material). Overall, the comparison with the four control simulations supports the results on the superagonist peptide, *i.e.* a higher structural stability and a stronger TCR association than the wild-type complex irrespective of the protonation state of the MHC acidic cluster.

**Supporting Tables**

| *Residue* | *crystal pK$_a$* | *pK$_a^1$* | *pK$_a^2$* |
|---|---|---|---|
| Glu11α | 7.4 | 6.9 | 8.3 |
| Asp66α | 11.1 | 11.4 | 8.7 |
| Asp11β | 13.7 | 8.8 | 8.5 |
| Asp30β | 5.7 | 6.1 | 8.6 |
| Asp37β | 12.1 | 7.4 | 6.0 |
| Asp57β | 4.1 | 3.7 | 4.0 |

**Table S1** pK$_a$ values as measured with PROPKA for the residues of the MHC acidic cluster, using structures of the original (unprotonated) wild-type complex. The table includes p$K_a$ values calculated for the X-ray structure, values for the structure after energy minimization (p$K_a^1$), and values for the structure after MD relaxation of solvent and sidechains (p$K_a^2$). Note that the values in the X-ray structure can be extremely high (*viz.* above 11, which is the range of values for a strong base) because of crystal packing.

| *Complex* | *Runs* | *Total sampling* |
|---|---|---|
| Wild-type | 10x1.08 μs | 10.8 μs |
| Aspp66 | 10x0.74 μs | 7.4 μs |
| Aspp11 | 10x0.83 μs | 8.3 μs |
| Aspp37 | 10x0.74 μs | 7.4 μs |

**Table S2** Single run and cumulative sampling time of tripartite complexes with different protonations; on Asp66α, Asp11β, or Asp37β. The additional "p" in the residue names stands for protonated. A sampling time between 700 ns and 800 ns was considered adequate for the control simulations. Note that despite the smaller sampling of each of the protonated systems a higher stability of the superagonist is observed.

**Supporting Figures**

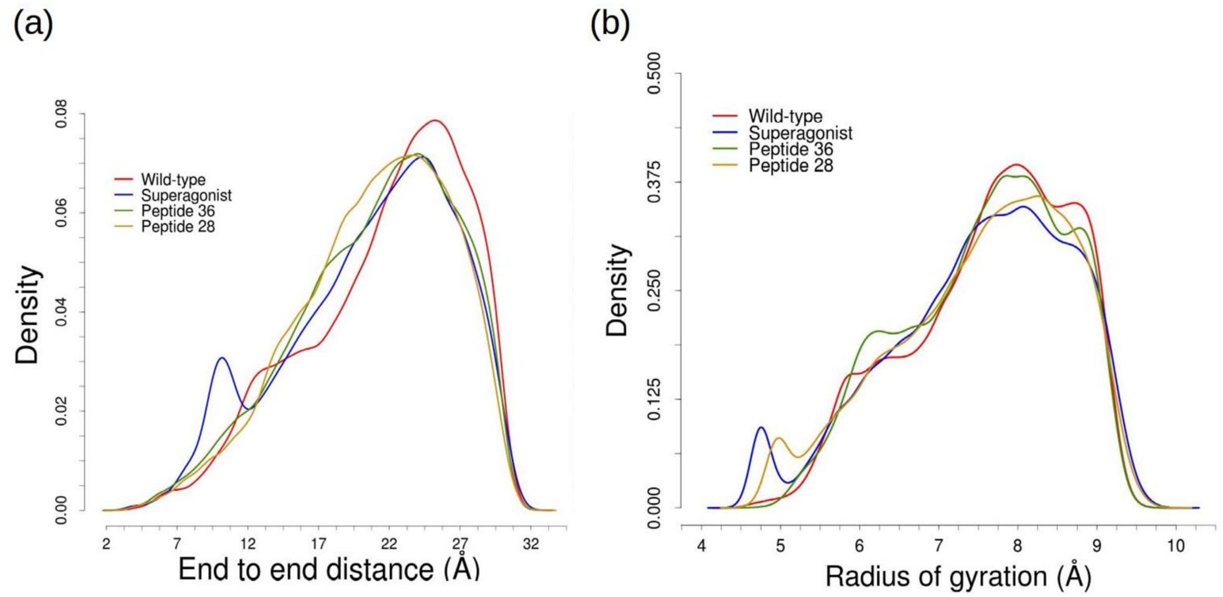


**Fig. S1** Density distribution of end-to-end distances (a) and radius of gyration (b) for the four free peptides. The wild-type peptide shows slightly higher values than the other peptides, indicating that it has the most extended conformation. The superagonist peptide shows a bimodal distribution, due to a compact state which is sampled mostly by a single copy (see Fig. S2 in the Supporting Material).

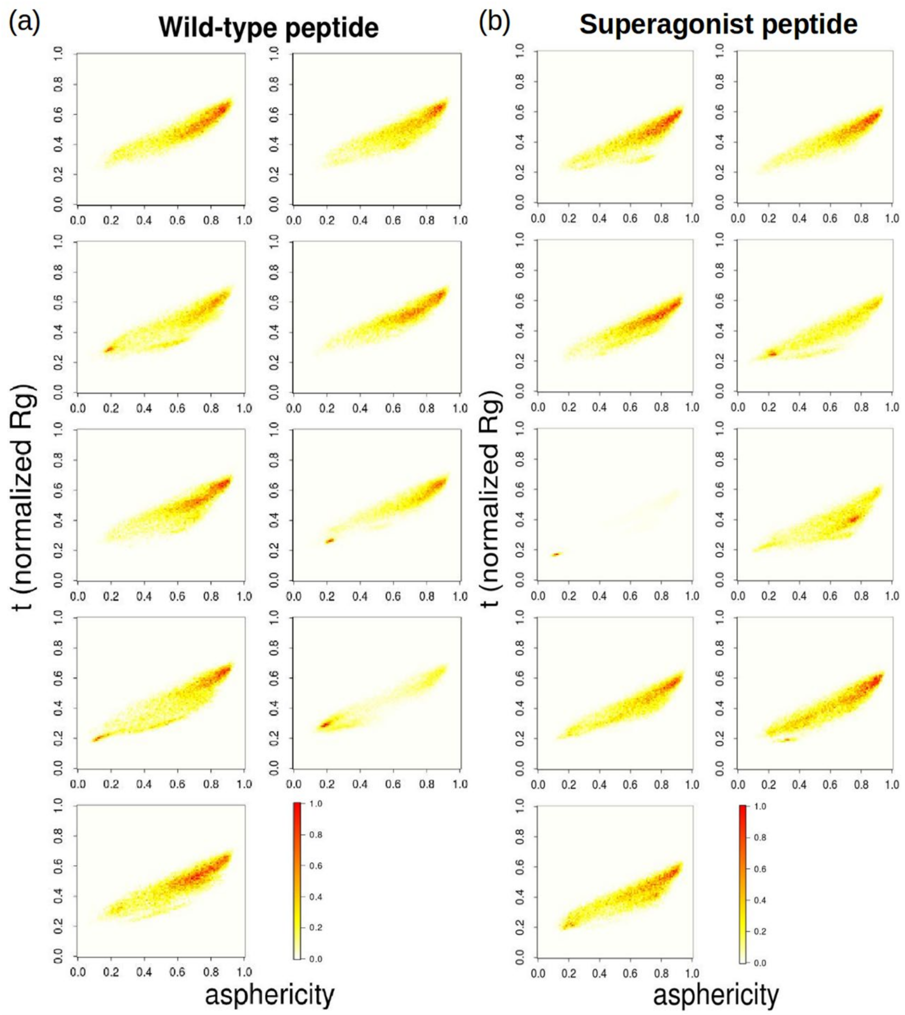


**Fig. S2** Statistical significance of the sampling of the peptide in the unbound state. The nine panels show the two-dimensional histograms of normalized radius of gyration and asphericity, plotted for every independent simulation run of wild-type (a) and superagonist (b). Both peptides exhibit one copy that samples almost uniquely a highly compact state. However, the individual runs are overall very similar in the regions sampled by the extended conformations. We conclude that the extended state shows statistical robustness.

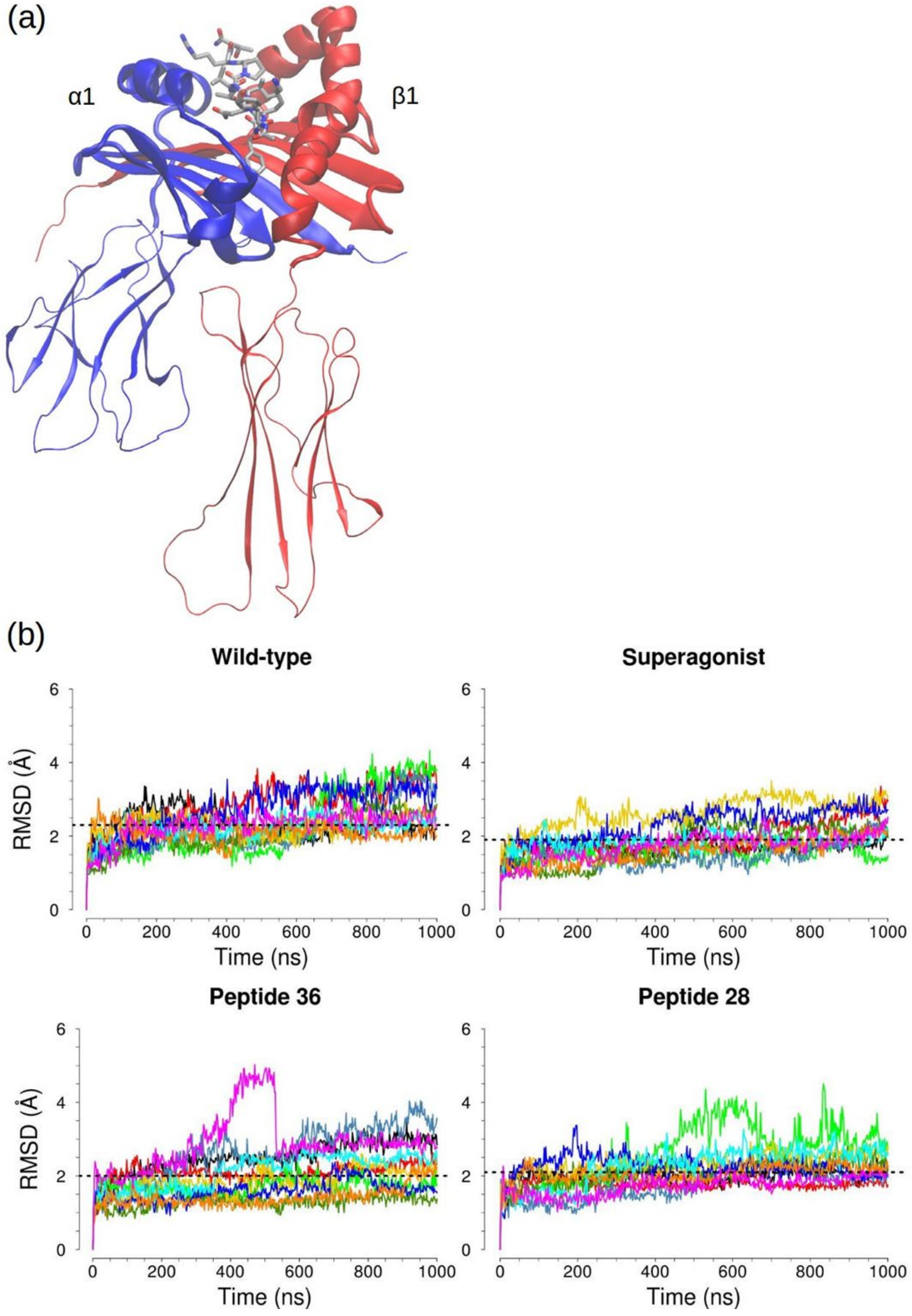


**Fig. S3** Visual representation and results of the RMSD analysis for the α1β1 helices after alignment on the MHCα1β1 domains. In (a) the MHC protein is shown in cartoon view, colored by chain, and the peptide in licorice (grey). Only residues in the recognition helices interact with TCR residues, while MHC helices and β-strands of the binding groove participate in peptide contacts. For the RMSD analysis, the MHCα1β1 domains (thick cartoon) were used for initial fitting and the RMSD was calculated only for the α1β1 helical residues (Cα atoms). (b) The overall behavior is similar for all systems, with the superagonist complex showing a slightly lower average (dashed line), *i.e.* a higher structural stability.

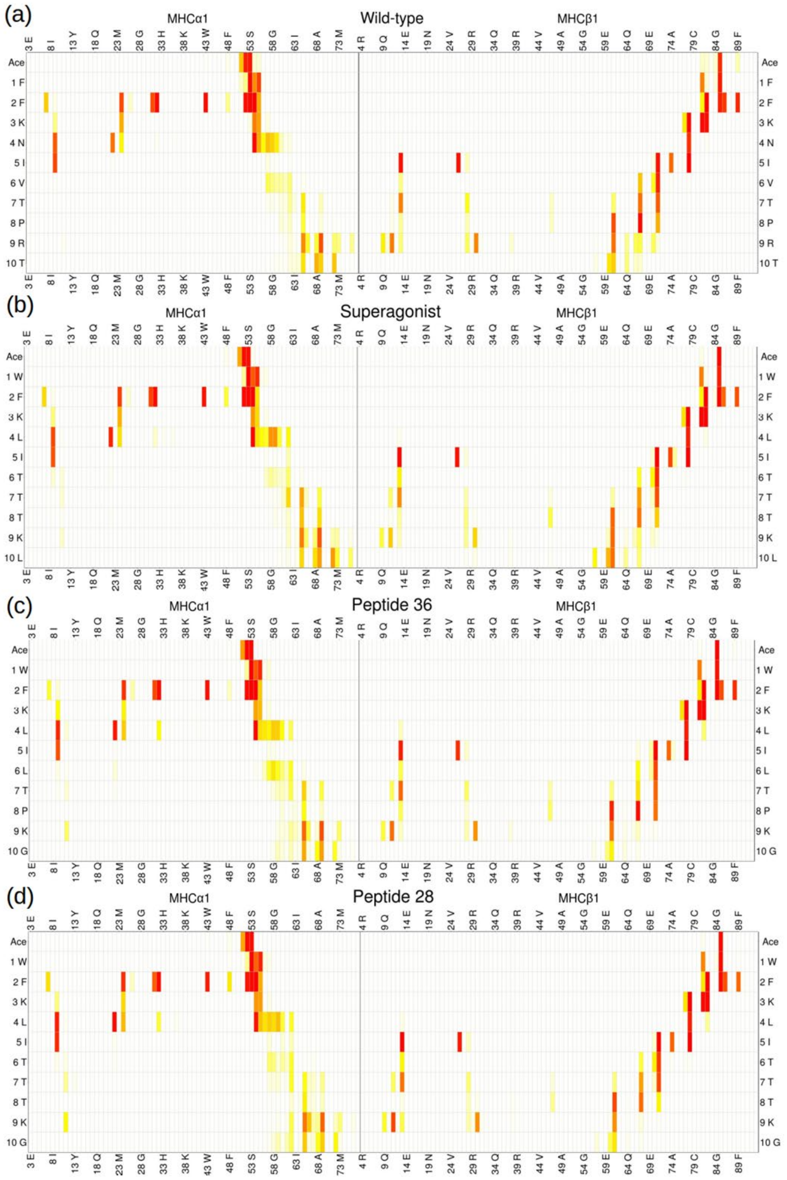


**Fig. S4** Contact maps for the pMHC runs, averaged over 10 copies per complex. The frequencies of contacts are shown for each complex as a heat map, with linear color scale as described and represented in Fig. 4. The pattern of contacts are very similar for all bipartite complexes and consistent with the reported binding motifs of MHC class II proteins.

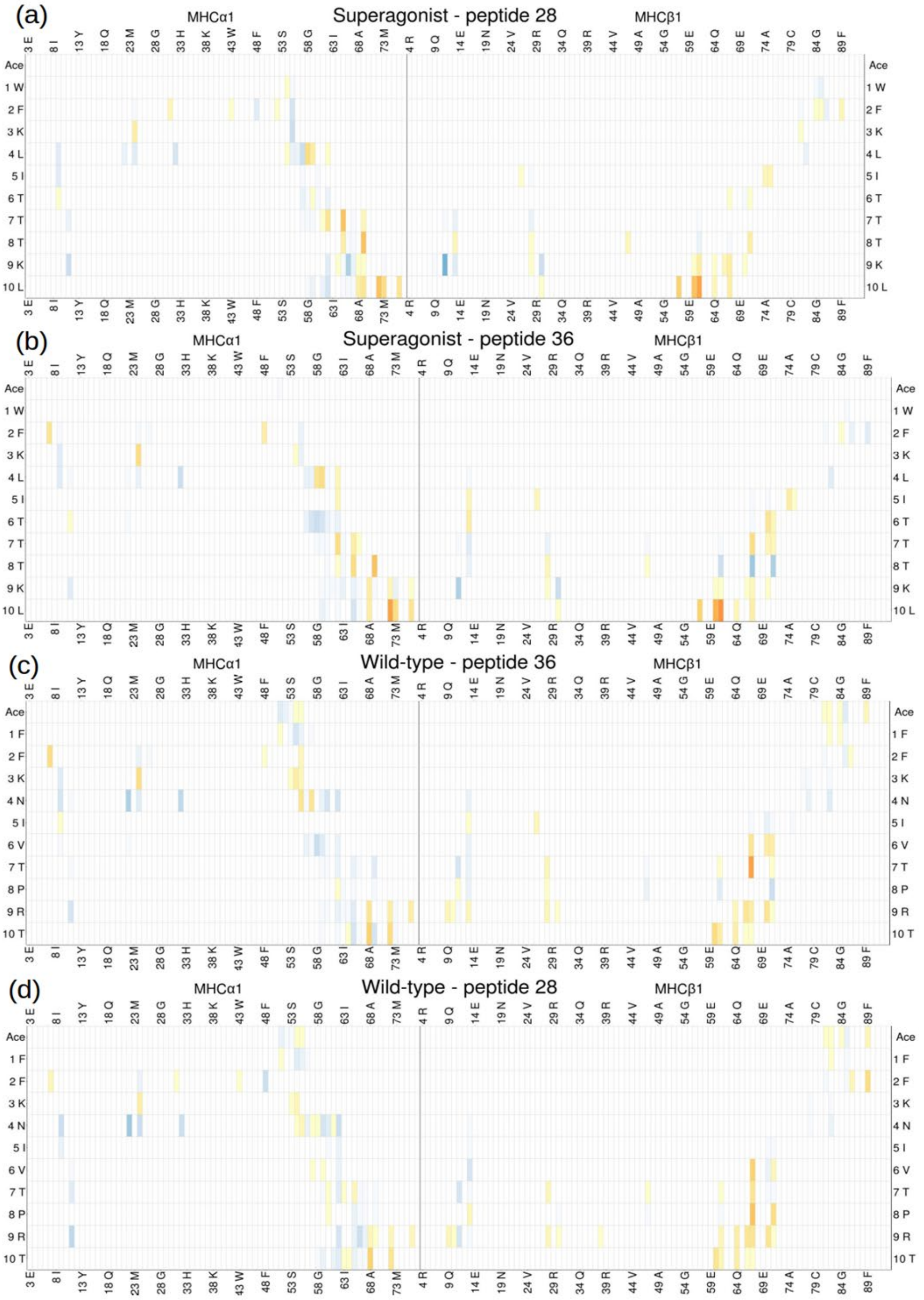


**Fig. S5** Difference contact maps. The linear color scale ranges from blue (higher frequency of contact for the second term) to red (higher frequency of contact for the first term), as described and represented in Fig. 4. The differences in contact frequencies are low, and localized mostly in the central and C-terminal regions of the peptide. Note the propagation of the stronger C-terminal contacts in the superagonist compared to peptide 28, despite a single residue (Leu10Gly) mutation.

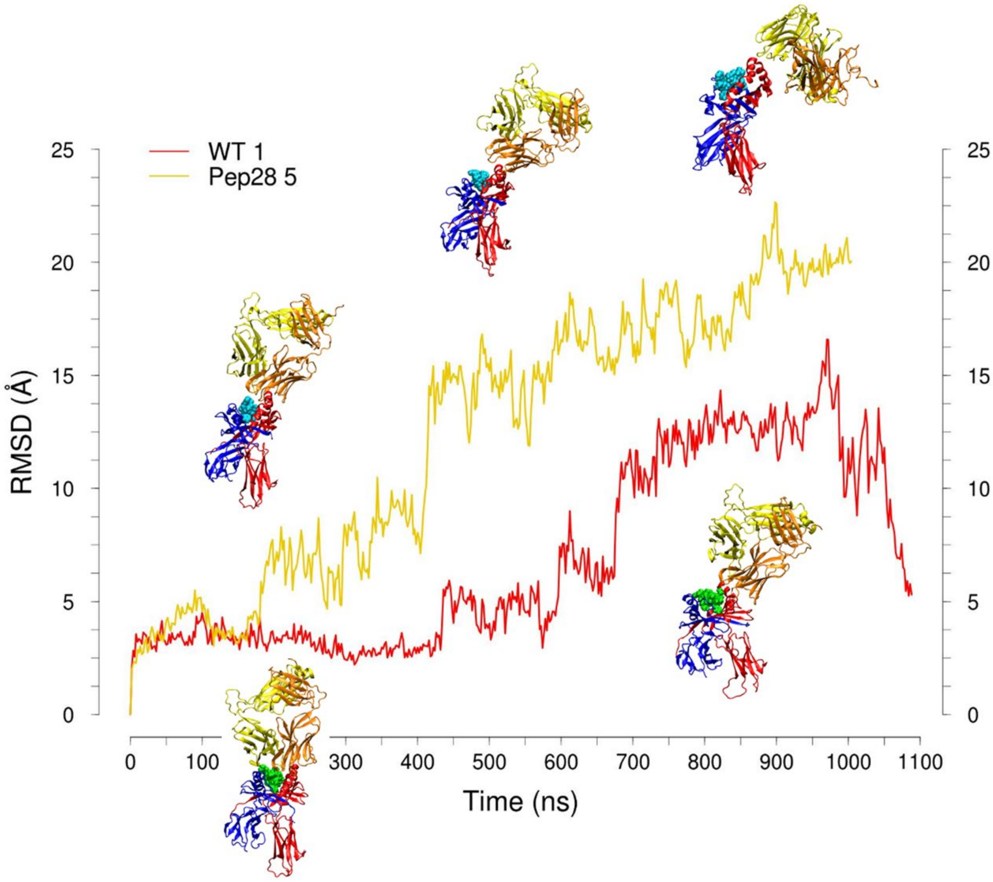


**Fig. S6** Time series of the RMSD of all Cα atoms for the runs with largest deviations in the wild-type and peptide 28 complexes. The wild-type copy (WT 1) has large shifts in the MHCβ2 domain (distant from the binding interface) and the TCRβ region loses contacts with the MHCα1 helix. The same contacts are also partially re-established at the end of the simulation, *i.e.* over 1μs (the wild-type runs were extended to verify the convergence of this copy to the sampled range). Peptide 28 copy 5 (Pep28 5) presents a gradual and complete loss of TCR contacts at the binding interface: initially the TCRβ chain loses its interactions, then the TCRα chain, and they both establish contacts on a lateral region of the MHCβ, leaving the peptide completely exposed to the solvent. We represent only the deviations for wild-type and peptide 28 because the highly deviating copies in superagonist and peptide 36 are due to persistent shifts in the MHCβ2 domain distant from the binding interface, without loss of TCR contacts on the MHC chains. As such, they do not influence the analysis and calculations limited to the binding interface, *viz.,* MHCα1β1 domains, TCR VαVβ regions and the peptide residues.

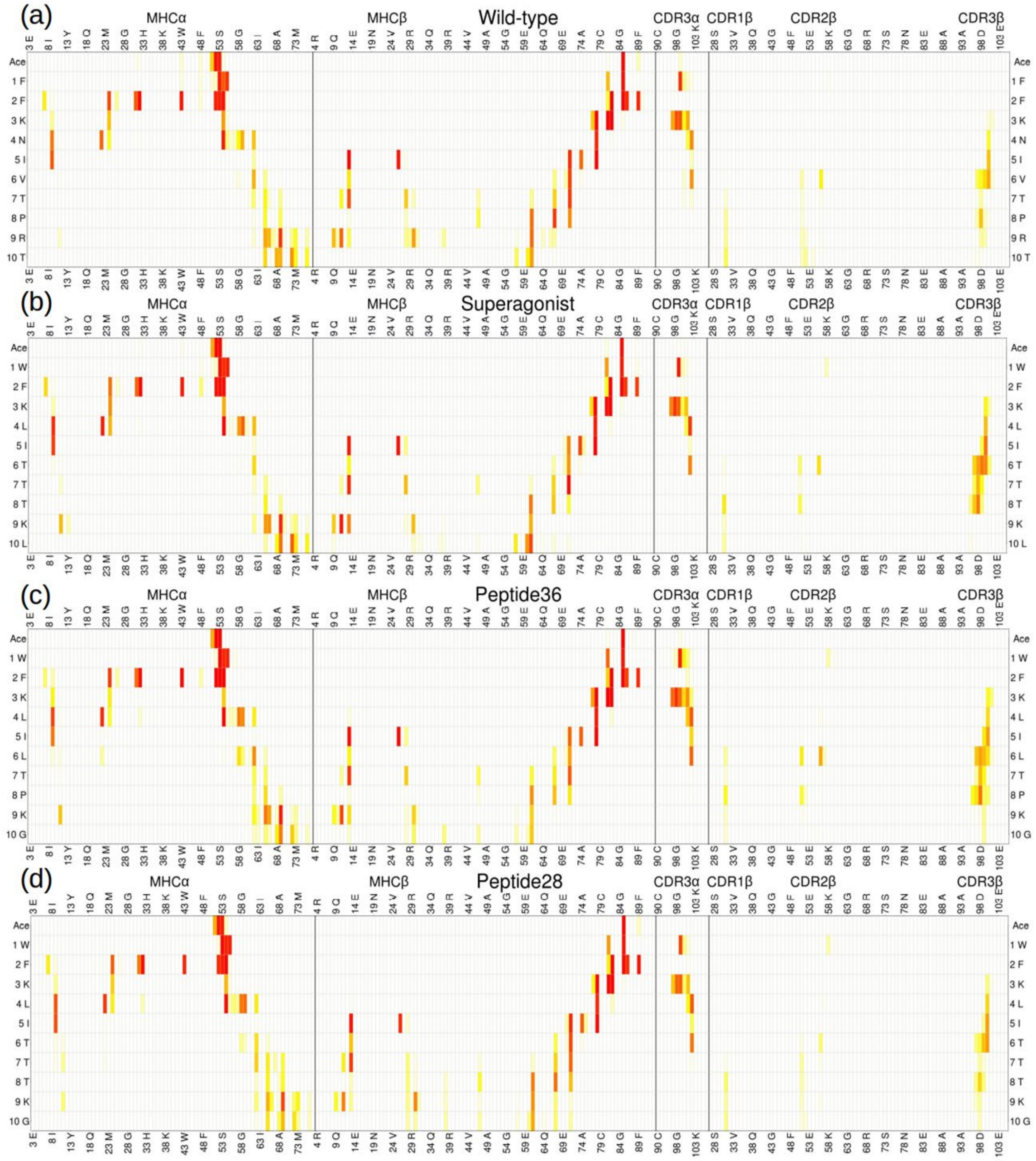


**Fig. S7** Contact maps of tripartite simulations, shown as a heat map with linear color scale as previously described. The patterns are similar for the four systems. The main differences in frequencies are localized in the peptide – TCR interactions, especially the contacts with the CDR3β loop.

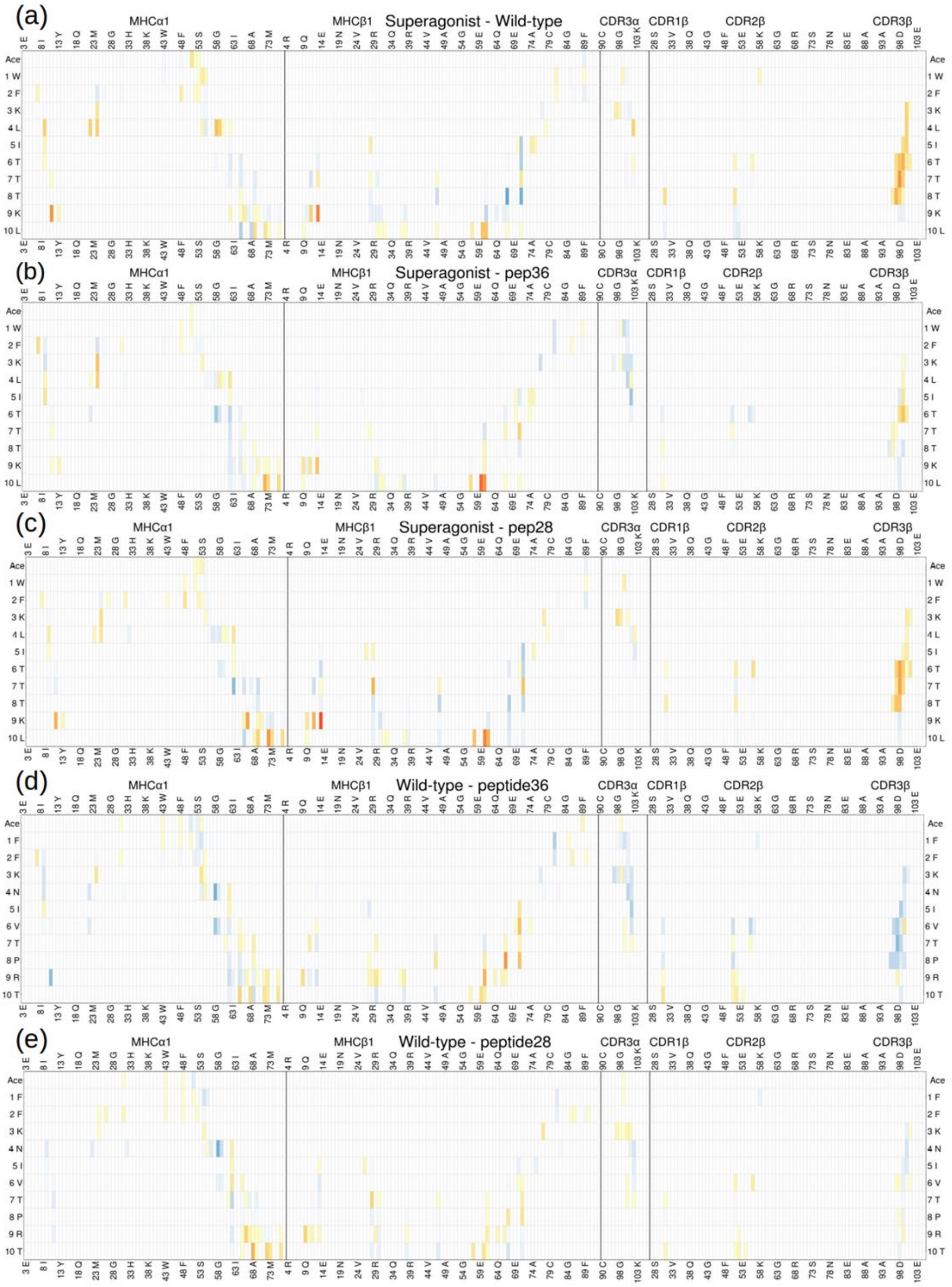


**Fig. S8** Difference contact maps. The linear color scale is the same as previously described for difference contact maps. The differences are most evident in the comparison with the superagonist peptide, *i.e.* in the three maps from the top, and involve mainly the central and C-terminal peptide residues.

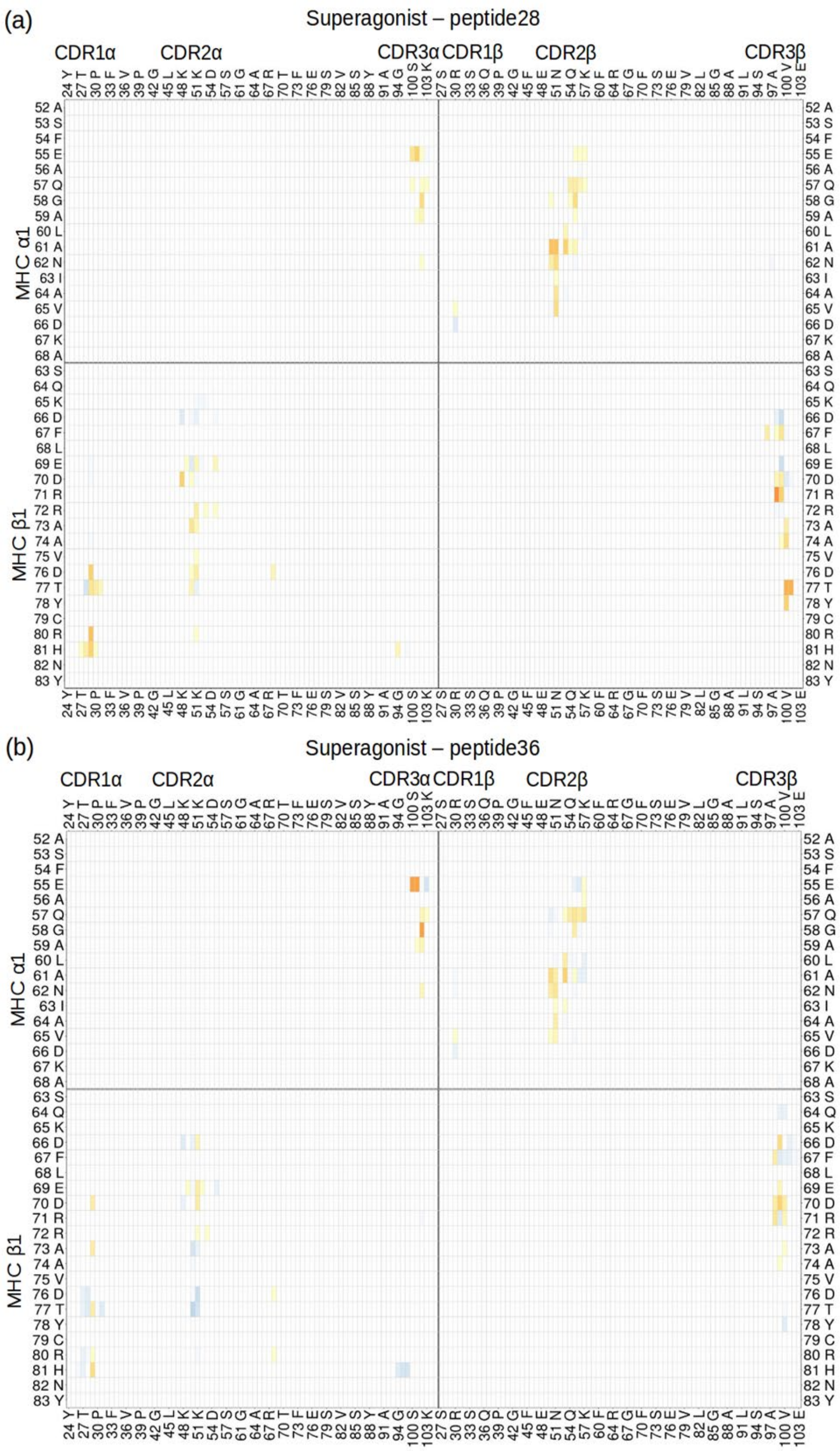


**Fig. S9** Difference contact maps. The plots show the same list of residues and linear color scale as Fig. 8b. The interactions of the CDR3β loop with MHCβ1 helix are stronger in the superagonist complex than in peptide 28 (a), which differ in the Gly10Leu mutation. The superagonist peptide also induces overall stronger MHC-TCR contacts with respect to peptide 36 (b).

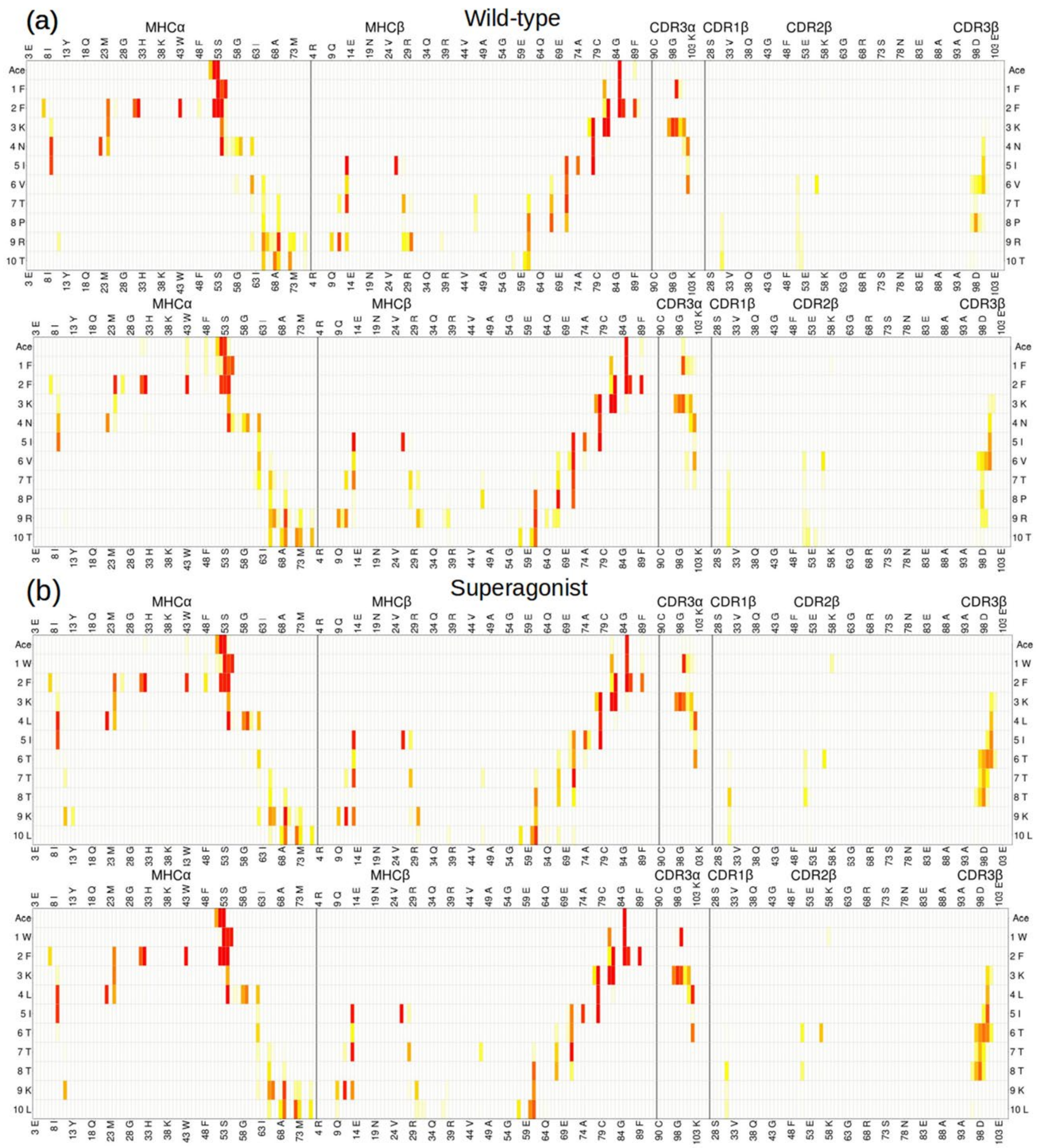


**Fig. S10** Block averaging of the contact maps involving peptide, MHC and TCR residues, with the same linear color scale as previous contact maps. We show here only the simulations for wild-type (a) and superagonist (b), and the same level of similarity was obtained for peptide 36 and peptide 28 complexes. All were divided in two equal sets of five randomly chosen trajectories, the contacts patterns were calculated and averaged over 5 runs. The high level of similarity between each set gives evidence for statistical robustness in our simulations.

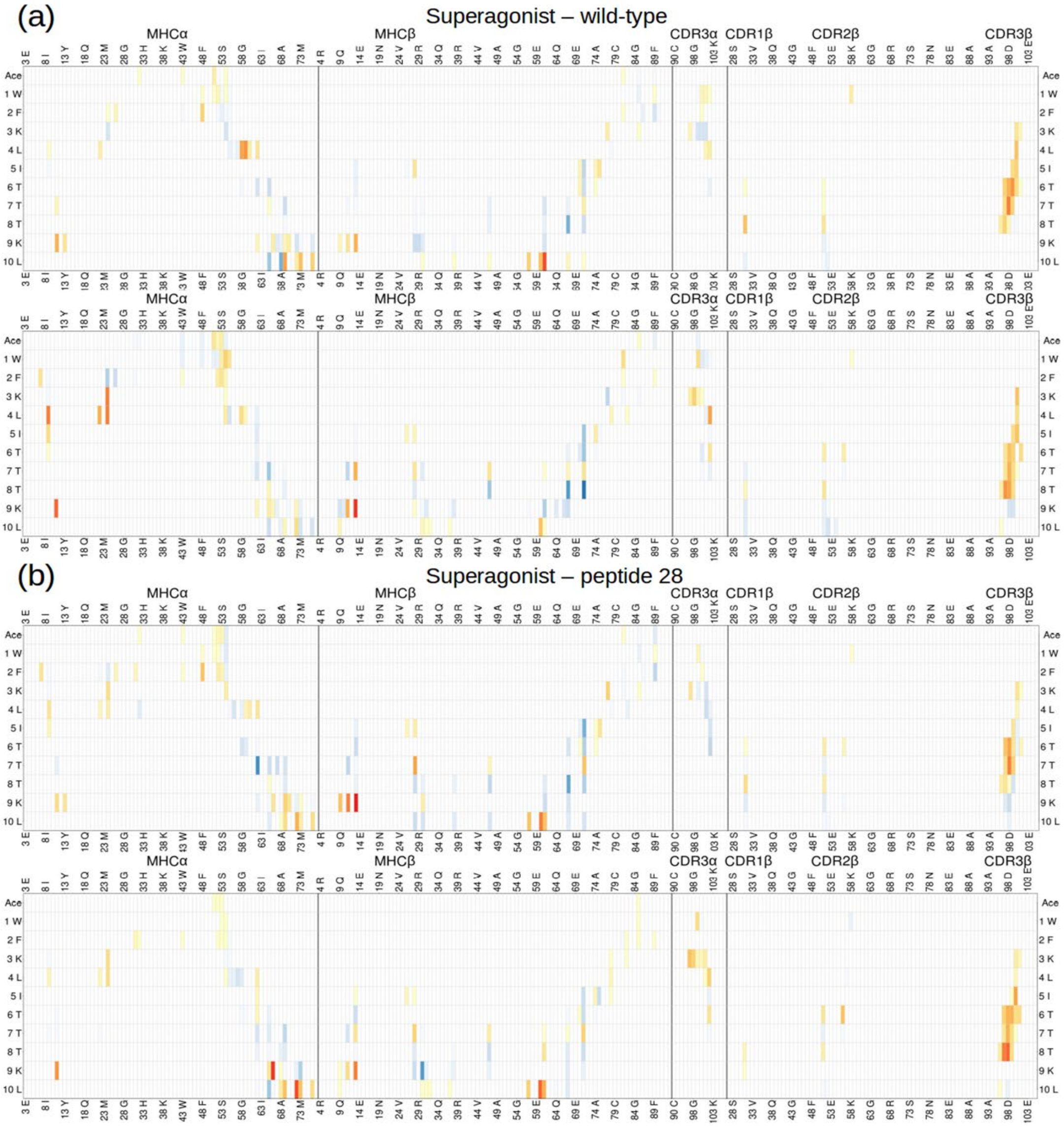


**Fig. S11** Block averaging of the difference contact maps involving peptide, MHC and TCR residues, with the same linear color scale as previous difference contact maps. The simulations indicated by both terms were divided in two equal sets of five randomly chosen trajectories (1 μs each) and the subtraction was applied for each set. We show here the block averages of comparisons we are most interested in, *i.e.* superagonist – wild-type comparison (a) and superagonist – peptide 28 comparison (b). The high level of similarity between each set gives evidence for statistical robustness.

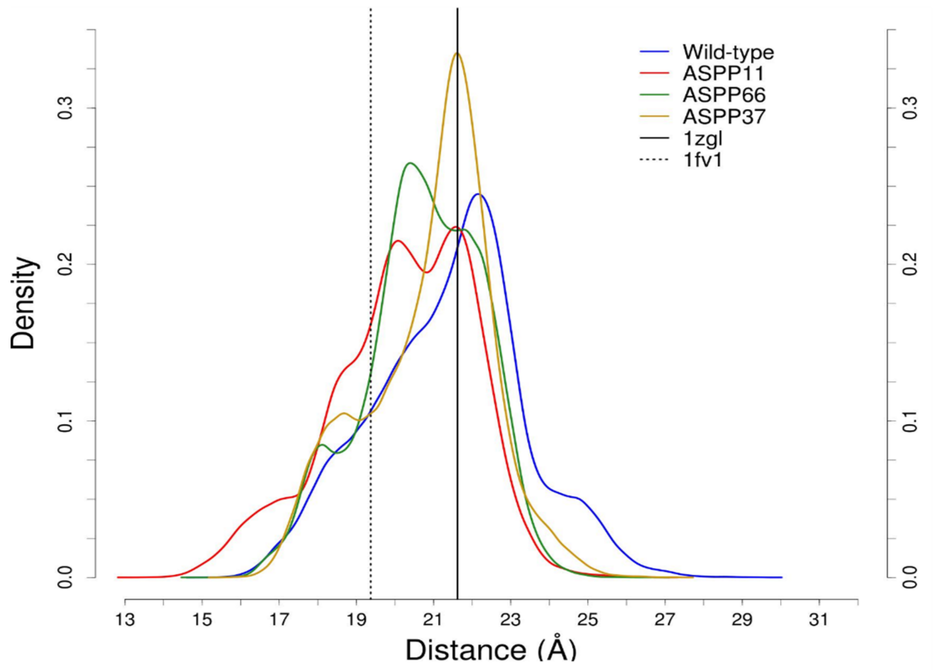


**Fig. S12** Density distributions for the D66β-V65α distances in four different MHC protonation states with the same wild-type peptide. The peaks and distributions are closer to the crystal structure value of the tripartite complex (vertical solid line) than the bipartite one (vertical dashed line).

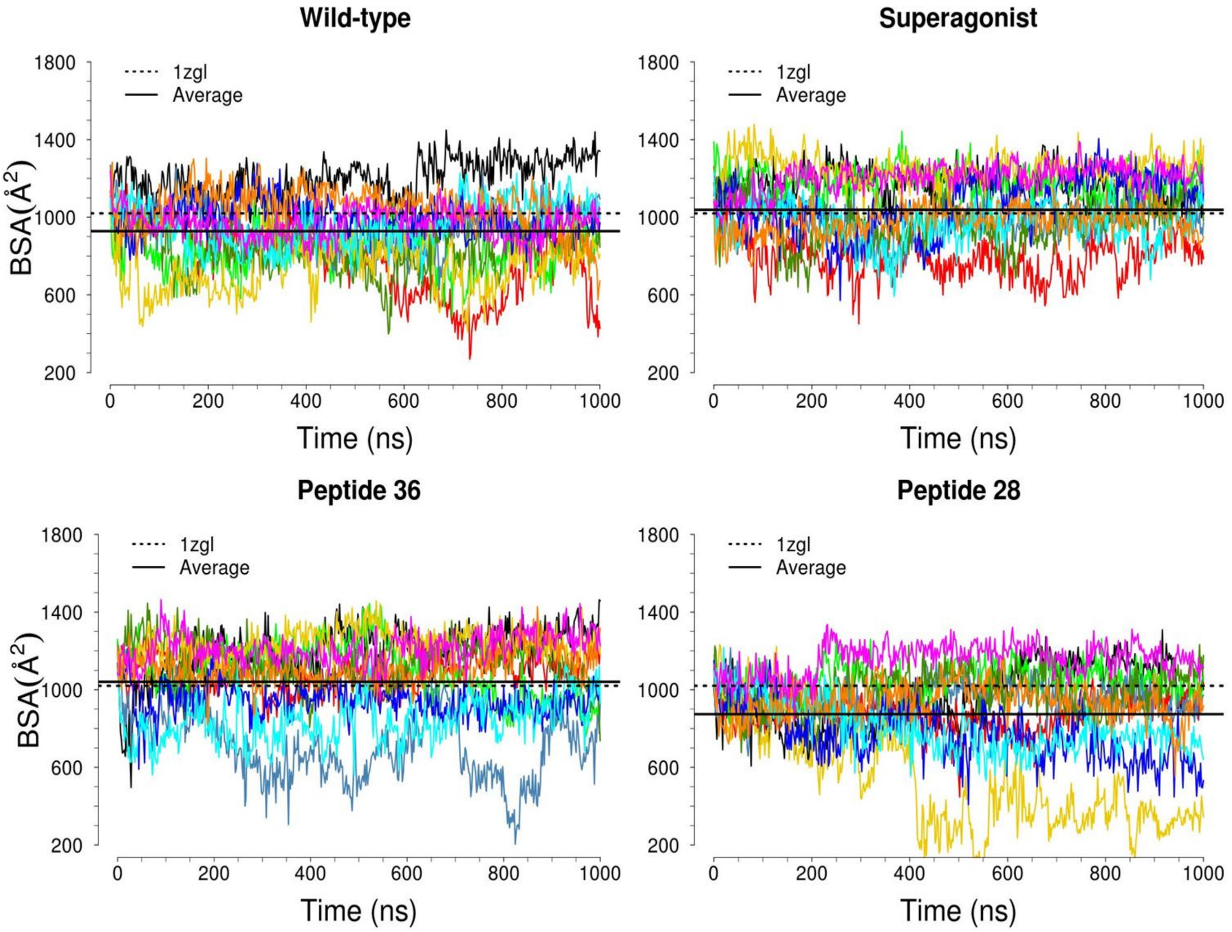


**Fig. S13** Temporal series of the buried surface area values for the four different peptide complexes. The superagonist shows higher values and the average (solid line) is close to the TCR surface buried in the crystal structure (dashed line). A similar result is obtained for peptide 36, whereas the wild-type and peptide 28 bury less TCR surface and are more distant from the crystal value. The superagonist shows the smallest fluctuations.

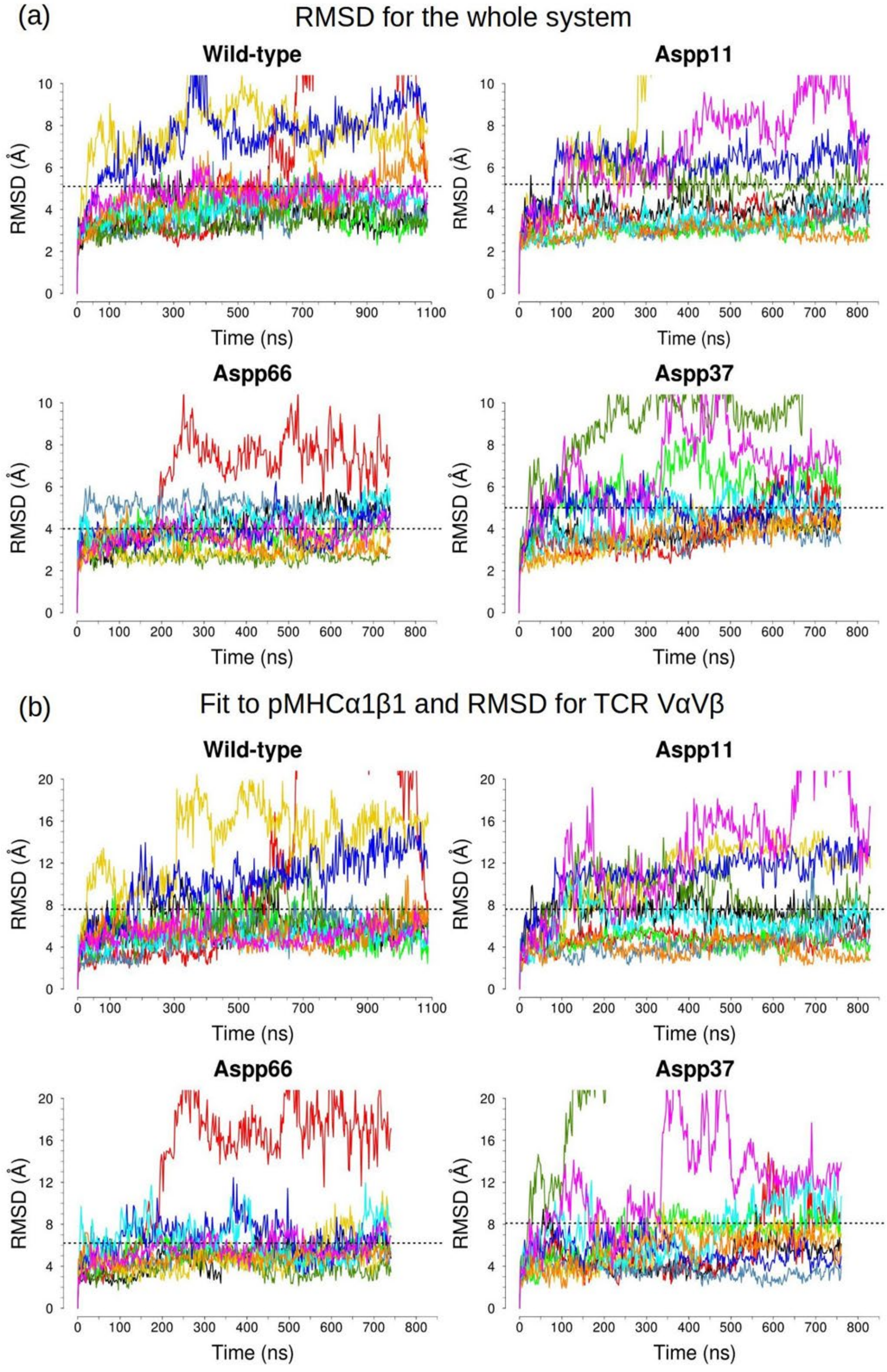


**Fig. S14** Temporal series of the RMSD calculations of all Cα atoms of the system (a) and the Cα of TCR VαVβ after fitting to MHCα1β1 (b), for the four protonation states. The time series show higher flexibility than the superagonist RMSD plots in Fig. 3b and Fig. 5c.

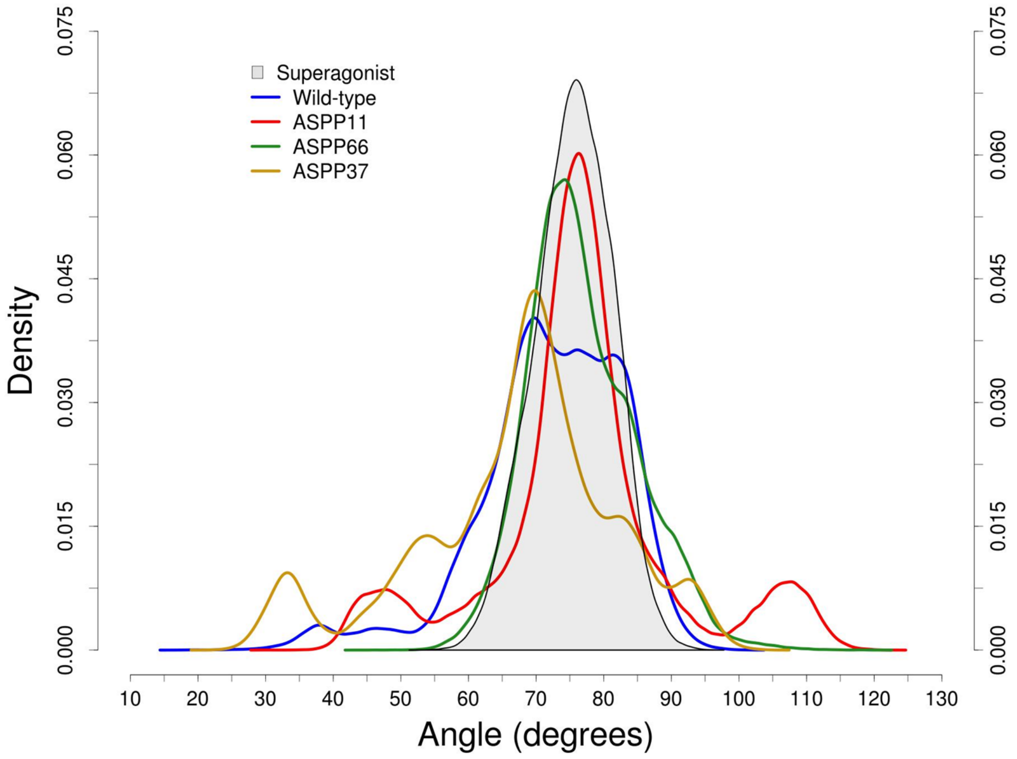


**Fig. S15** Density distributions of the orientation angle of TCR chains with respect to the pMHC surface for the protonated controls and the superagonist complex (same distribution as in Fig. 6c). The results show a greater variability for all the protonation states of wild-type with respect to the superagonist which maintains the TCR position very stable over the pMHC surface.

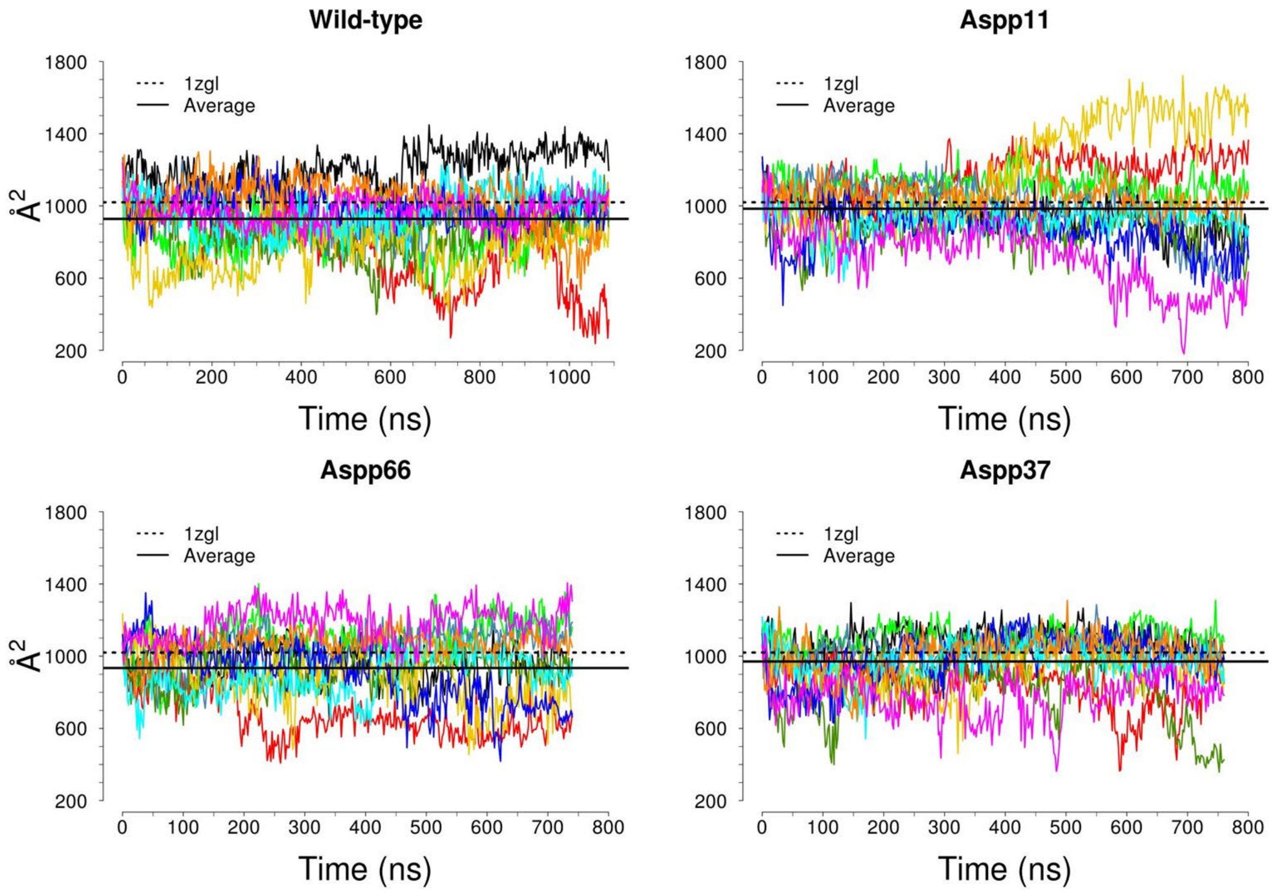


**Fig. S16** Temporal series of the buried surface area (BSA) of TCR chains for the four protonation states of the MHC. The results show that the protonation states are similar between themselves, except for larger deviations in the Aspp11 complex. Overall, the protonated complexes show slightly higher deviations than the superagonist (BSA plot in Fig. S13).
